# Substrate-triggered position-switching of TatA and TatB is an essential step in the *Escherichia coli* Tat protein export pathway

**DOI:** 10.1101/113985

**Authors:** Johann Habersetzer, Kristoffer Moore, Jon Cherry, Grant Buchanan, Phillip Stansfeld, Tracy Palmer

## Abstract

The twin arginine protein transport (Tat) machinery mediates the translocation of folded proteins across the cytoplasmic membrane of prokaryotes and the thylakoid membrane of plant chloroplasts. The *Escherichia coli* Tat system comprises TatC and two additional sequence-related proteins, TatA and TatB. Here we use disulfide crosslinking and molecular modelling to show there are two binding sites for TatA/B proteins on TatC. TatA and TatB are each able to occupy both sites if they are the only TatA/B protein present. However, under resting conditions the sites are differentially occupied with TatB occupying the ‘polar cluster’ site while TatA binds adjacently at the TatC transmembrane helix 6 binding site. When the Tat system is activated by the overproduction of a substrate, TatA and TatB switch their binding sites. We propose that this substrate-triggered positional exchange is a key step in the assembly of an active Tat translocase.

## Introduction

The twin-arginine protein transport (Tat) pathway operates in parallel with the general secretory (Sec) pathway to export proteins across the cytoplasmic membrane of bacteria and the thylakoid membrane of plant chloroplasts. Substrates are targeted to the Tat pathway by N-terminal signal peptides containing a conserved twin-arginine motif and are transported across the membrane in a folded state driven by the protonmotive force (Berks, 2015;Cline, 2015;Palmer and Berks, 2012).

The Tat machinery comprises membrane proteins from the TatA and TatC families. TatA family proteins are monotopic with an N-out transmembrane helix at their N-terminus, followed by a cytoplasmically-located amphipathic helix (Aldridge et al., 2012; Koch et al., 2012). Most Gram-negative bacteria, and plant thylakoids, have two functionally distinguishable TatA paralogues, termed TatA and TatB in bacteria (Tha4 and Hcf106 in plant thylakoids), that have distinct roles in Tat transport (e.g.Cline and Mori, 2001; Sargent et al., 1999). TatC is the core component of the Tat system, and forms a scaffold for the dynamic assembly of Tat complexes during protein translocation (Ramasamy et al., 2013; Rollauer et al., 2012).

Tat transport is initiated by binding of the signal peptide of a Tat substrate to the Tat(A)BC receptor complex. This complex, which contains several copies of TatB and TatC, is multivalent and appears to function as an obligate oligomer (Bolhuis et al., 2001; Cleon et al., 2015; Ma and Cline, 2010; Tarry et al., 2009). Although the Tat(A)BC complex is stable and can interact with substrates in the absence of TatA (Behrendt and Bruser, 2014; Tarry et al., 2009), it is likely that *in vivo* some TatA constitutively associates with this complex, most likely in an equimolar ratio with TatB and TatC (Alcock et al., 2016; Aldridge et al., 2014; Bolhuis et al., 2001; Zoufaly et al., 2012).

The twin arginine motif of the signal peptide is recognised by the cytoplasmic surface of TatC (Gerard and Cline, 2007; Rollauer et al., 2012). The signal peptide can also bind more deeply within the receptor complex, contacting residues in the transmembrane helix (TM) of TatB and towards the periplasmic end of TatC TM5 (Alami et al., 2003; Blummel et al., 2015; Gerard and Cline, 2007). Following substrate binding, additional TatA protomers are recruited to the receptor complex dependent on the protonmotive force (Alami et al., 2003; Alcock et al., 2013; Aldridge et al., 2014; Dabney-Smith and Cline, 2009; Dabney-Smith et al., 2006; Mori and Cline, 2002; Rose et al., 2013). According to current models, the assembled TatA oligomer facilitates substrate translocation across the membrane either through formation of a size-variable channel or by promoting localised membrane weakening and transient bilayer disruption (see (Berks, 2015; Cline, 2015) for recent reviews).

Although high resolution structural information is available for TatA, TatB and TatC (Hu et al., 2010; Ramasamy et al., 2013; Rodriguez et al., 2013; Rollauer et al., 2012; Zhang et al., 2014a; Zhang et al., 2014b), to date Tat complexes have only been visualised at low resolution (Oates et al., 2003; Sargent et al., 2001; Tarry et al., 2009). Site-specific crosslinking has been used to map interaction interfaces between Tat components, giving results consistent with a potential binding site for TatB and/or TatA being located at TM5/TM6 of TatC (Aldridge et al., 2014; Blummel et al., 2015; Kneuper et al., 2012; Rollauer et al., 2012). Recently, co-evolution analysis independently predicted the location a TatA/TatB binding site at TM5/6 of TatC, pointing to a polar cluster of amino acids in *E. coli* TatC (M205, T208 and Q215) forming likely contacts with a polar side chain in TatA and TatB (Q8 or E8, respectively, in the *E. coli* proteins) (Alcock et al., 2016). Site-directed mutagenesis confirmed the importance of these polar residues for TatA and TatB interaction with TatC and supported the generation of molecular-level resolution models of TatAC and TatBC complexes. From this it was inferred that TatA and TatB share a common binding site on TatC, which is differentially occupied by these proteins at different stages of Tat transport (Alcock et al., 2016).

In this work we have undertaken an *in vivo* disulfide crosslinking study to explore the interaction of TatC with TatA and TatB in the absence of a bound substrate and when a substrate is likely to be bound. Our studies identify two binding sites for each protein. The first of these, identified by Alcock et al (2016) is occupied by TatB under resting conditions, with TatA occupying a second binding site primarily located at TatC TM6. We go on to show that in the presence of over-expressed Tat substrate TatA and TatB move positions to occupy each other’s binding sites. We propose that signal peptide-triggered position switching of TatA and TatB is a critical step in driving the assembly of an active Tat translocase.

## Results

### The TatB TM is positioned close to TM5 of TatC at the polar cluster site under resting conditions

Prior studies analyzing disulfide crosslinking between *E. coli* TatB and TatC have used isolated membrane fractions harboring elevated copies of Tat components produced from multicopy plasmids. Under these conditions an initial contact site between TatB^L9C^ and TatC^M205C^ was identified (Kneuper et al., 2012), which was subsequently extended to reveal further contacts between Cys residues introduced into the TM of TatB and into TM5 of TatC (Rollauer et al., 2012). To explore whether the same contact sites were detectable *in vivo,* we developed a protocol for disulfide crosslinking in intact cells using the TatB^L9C^ -TatC^M205C^ crosslink. In these experiments, the Cys-substituted variants of TatB and TatC were produced from the low copy number plasmid p101C*BC, which expresses *tatBC* at approximately chromosomal level (Alcock et al., 2013), in a strain lacking chromosomal *tatBC.* An initial titration with the oxidant copper phenanthroline (CuP) revealed that a TatBC crosslink was detectable when CuP was used at 1.2 and 1.8mM (Fig 1A). We also noted that TatB and TatC homodimers were formed through the introduced Cys residues after incubation with CuP, as reported previously (Kneuper et al., 2012; Rollauer et al., 2012). Next, using 1.8mM CuP, we undertook a time course from 1 -15 min and examined the formation of the TatBC heterodimer and the survival of cells during this period. Fig 1B shows a TatBC heterodimer was detected at all time points, including the earliest time point tested, however, incubation times with CuP in excess of one minute saw a significant reduction in the recovery of cells (Fig 1C). We therefore chose to use a 1 min incubation with 1.8mM CuP for all subsequent crosslinking analysis.

**Figure 1.**
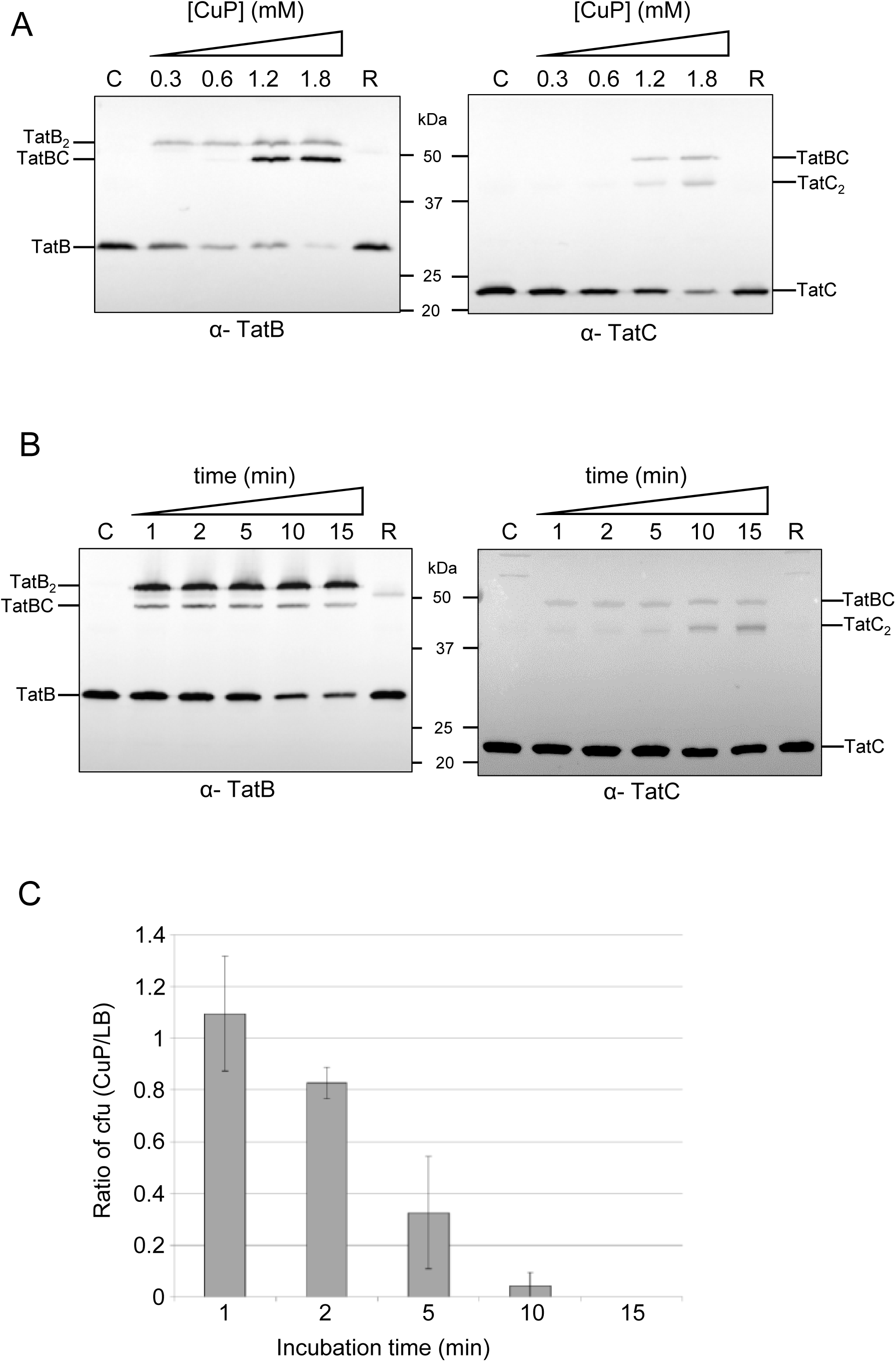
Development of an *in vivo* disulfide crosslinking protocol. Cells of strain MC4100∆BC (*∆tatBC*) harboring plasmid p101C*BC producing TatB^**L9C**^ alongside TatC^**M205C**^ were incubated with either LB medium (Control, C), or LB supplemented with 10 mM DTT (reduced; R) or **A.** the indicated concentrations of CuP for 15 min or **B.** with 1.8 mM CuP for 1-15 min. The reaction was quenched by addition of 8 mM NEM/12 mM EDTA, membranes were prepared and proteins were separated by SDS-PAGE (10% polyacrylamide). Crosslinked products were visualized by immunoblotting using anti-TatB_FL_ (left panel) or anti-TatC (right panel) antibodies. **C.** Aliquots of cells from the oxidized and control samples in part B were spread on LB plates containing chloramphenicol and the number of colonies enumerated following growth at 37°C for 24 hours. The y-axis shows the ratio of the number of colony forming units (cfu) obtained after incubation with 1.8mM CuP compared to the number after incubation in LB medium only. *n* = 3 biological replicates, error bars are ± SD.

Next we introduced Cys residues into a scanning region of TatC from residue 205 in TM5, through the periplasmic P3 loop as far as residue 216 in TM6 (Fig 2A). Fig S1A shows that when each of these TatC Cys substitutions was co-produced with TatB^L9C^, cells were able to grow in the presence of 2% SDS. This indicates successful export of Tat substrates AmiA and AmiC (Ize et al., 2003) and therefore that the Cys substitutions did not abolish Tat transport activity. Following incubation of cells producing each of these variants with CuP, a TatBC heterodimer was primarily detected between TatB^L9C^ and TatC^M205C^ (Fig 2B,C). A faint TatBC heterodimer band was also seen between TatB^L9C^ and TatC^L206C^, and a fainter one between TatB^L9C^ and TatC^F213C^ that was only detected with the anti-TatB antibody (indicated with asterisks on Fig 2B). It should be noted that the TatB antiserum used in this scanning experiment is a polyclonal anti-peptide antibody that primarily recognizes the C-terminal 15 amino acids of TatB and detects the TatB homodimer as a doublet band (Fig S2) for reasons that are unclear.

**Figure 2.**
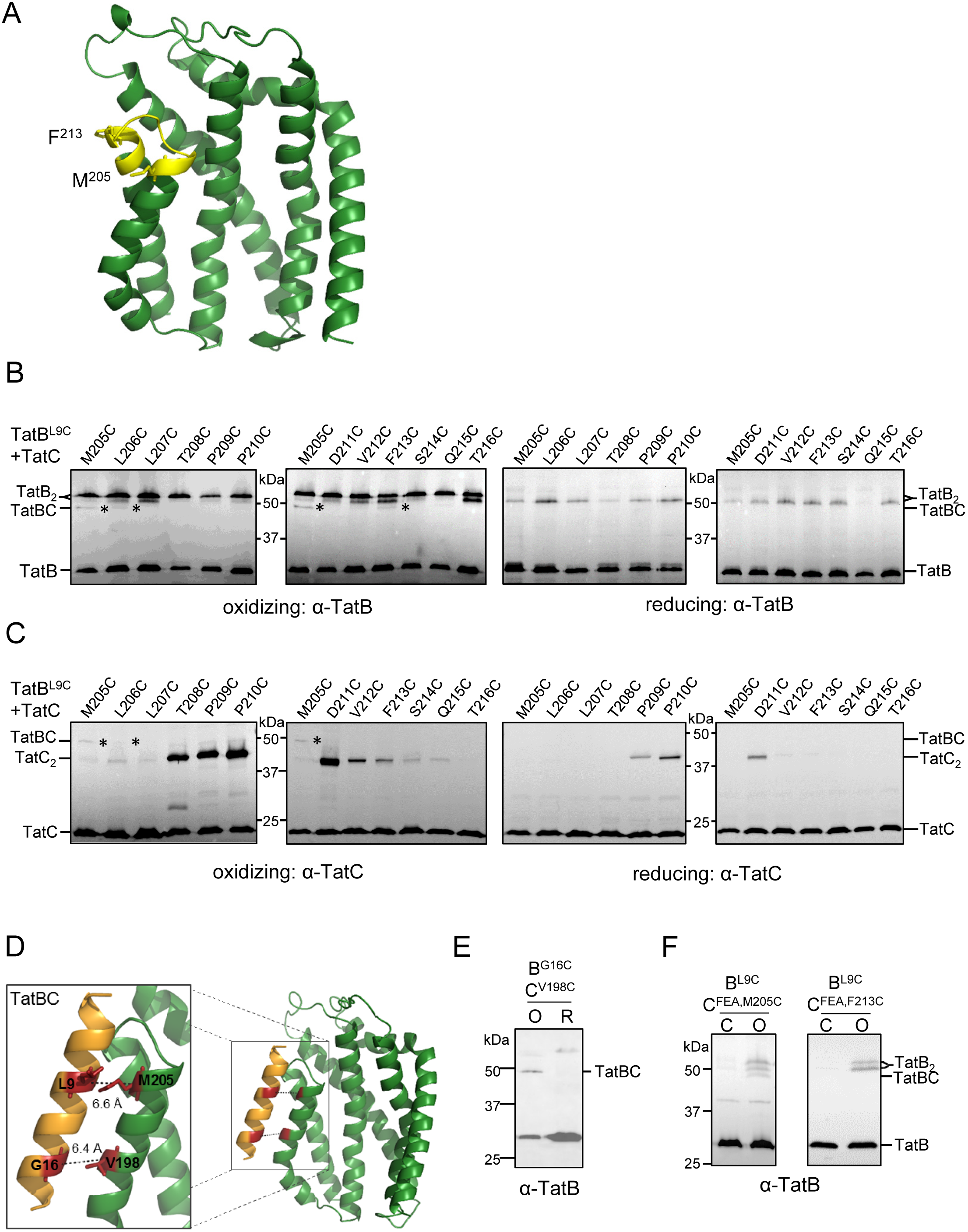
TatB^L9C^ interacts with TatC^M205C^ *in vivo.* A. Homology model of *E. coli* TatC showing positions of the residues used for disulfide crosslinking analysis in cyan. The side-chains of M205 and F213 are indicated. **B** and **C.** Western blot analysis (separated on 10% polyacrylamide gels) of membranes from *E. coli* strain MC4100∆BC producing TatB^**L9C**^ alongside the indicated Cys substitutions in TatC (from plasmid p101C*BC) following exposure of whole cells to 1.8mM CuP (oxidizing) or 10 mM DTT (reducing) for 1 min. Crosslinked products were visualized by immunoblotting using B. an anti-TatB peptide antibody or C. an anti-TatC antibody. The asterisks indicate likely TatBC crosslinks. **D.** Structural model of TatB interacting with TatC at the polar cluster site (adapted from Alcock et al., 2016). The backbone distances between TatB^L9^/TatC^M205^ and TatB^G16^/TatC^V198^ are shown. **E.** Whole cells of strain MC4100∆BC producing TatB^**L9C**^ alongside TatC^F94A,E103,/M205C^ or TatC^F94A/E103A/F213C^ from plasmid p101C*BC (annotated TatC^FEA,M205C^ or TatC^FEA,F213C^, respectively) were left untreated (C) or incubated for 1 min with 1.8mM CuP (O) as indicated. Following membrane preparation, crosslinks were detected with an anti-TatB peptide antibody.

Alcock et al. (2016) identified a binding site for TatA/TatB close to the polar cluster of residues M205, T208, Q215 in TatC. Molecular dynamics simulations indicated that TatB E8 may hydrogen bond with both of T208 and Q215 when bound at this site. We were unable to explore this directly by disulfide crosslinking since a Cys substitution at TatB^E8^ abolished Tat activity when expressed from plasmid p101C^*^BC (Fig S1A). This is consistent with the loss of activity noted for a TatB^E8A^ substitution, which resulted in destabilization of the TatB-TatC interaction (Alcock et al., 2016). However, molecular modelling indicates that when TatB interacts with TatC via the polar cluster, L9 of TatB is positioned within 6.6Å of TatC^M205^ (backbone distances; Fig 2D). We therefore conclude that the disulfide crosslink formed between TatB^L9C^ and TatC^M205C^ arises from interaction of TatB at the TatC polar cluster site. To confirm this we undertook disulfide crosslinking between TatB^G16C^ and TatC^V198C^ (Fig 2D) which are one of the most highly co-varying pair of residues at the polar cluster site (Alcock et al., 2016). Fig 2E shows that, as expected, a crosslink is formed between these two cysteine residues when cells were oxidized. These results give full support to the binding mode of TatB described previously (Alcock et al., 2016).

Experiments using a variant of TatC that is unable to bind signal peptides (TatC^F94A,E103A^) led to the conclusion that the interaction of TatB at the TatC polar cluster site occurred when the Tat system was at rest (Alcock et al., 2016). Fig 2F shows that in agreement with this, introduction of these same TatC substitutions did not abolish the TatB^L9C^ and TatC^M205C^ crosslink. Thus TatB occupies the polar cluster binding site under resting conditions *in vivo.*

### In the resting Tat system TatA interacts at a distinct site on TatC close to TM6

Next we used a similar approach to determine whether we could detect *in vivo* interactions between TatA and TatC. Initially, Cys-substituted variants of TatA and TatC were produced alongside TatB under control of the *lac* promoter from plasmid pQE60 and expressed in strain DADE-P (Lee et al., 2006) that lacks chromosomally encoded *tatABC/tatE* and which harbors the *pcnB1* allele to limit plasmid copy number to 1-2 per cell (Lopilato et al., 1986). Cys substitutions were introduced at L9, L10 and I11 of TatA and these were tested with the same Cys-scanning region from residues 205 - 216 of TatC.

First we confirmed that Tat activity was not abolished following introduction of any of these substitutions by showing that each pair of Cys-substituted proteins was able to support growth of DADE-P in the presence of 2% SDS (Fig S1B,C). Subsequently we undertook crosslinking analysis *in vivo* using the same protocol as that used for TatB-TatC crosslinking. Under the conditions tested, no crosslinks were detected between TatA and the TatC polar-cluster residue M205C (Fig 3A, Fig S3), indicating that TatA is not present at this site. Instead, a band of the expected size for a TatA-TatC heterodimer was detected under oxidizing conditions in cells co-producing TatA^L9C^ and TatC^F213C^. This was confirmed as a crosslink between TatC and TatA since it was also cross-reactive with an anti-TatA antibody (Fig 3B). A similar band was also detectable under oxidizing conditions when TatA^L9C^ and TatC^F213C^ were produced alongside wild type TatB at much lower levels from plasmid pTAT101 (Fig 3C). Scanning analysis using TatA^L10C^ revealed a faint crosslink with TatC^V212C^ following oxidation (Fig S3A), and TatAI11C gave detectable crosslinks with TatC^V212C^ and TatC^F213C^ (Fig S3B).

**Figure 3.**
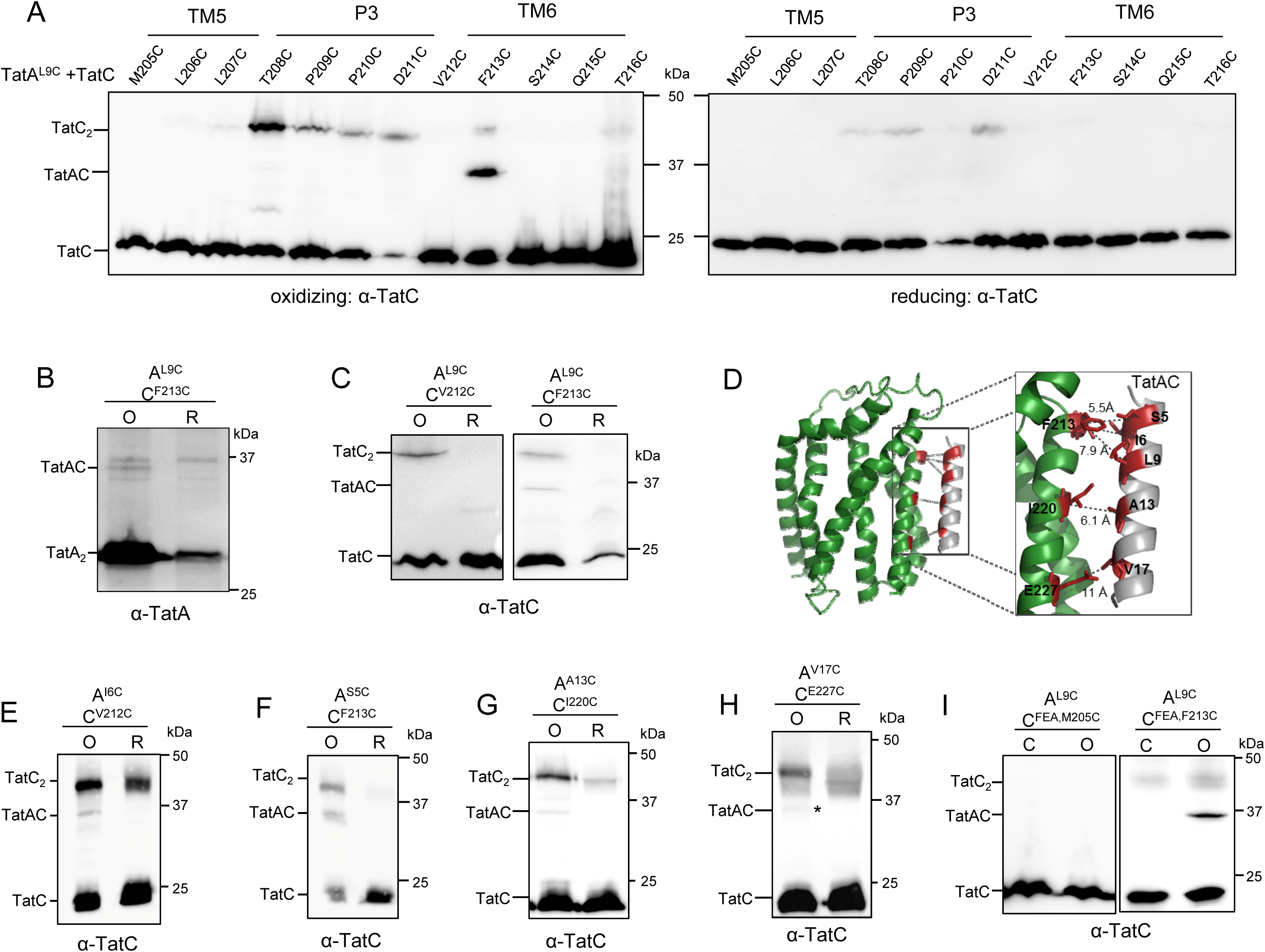
TatA^L9C^ interacts with TatC^F213C^ *in vivo.* A, E, G and H. Western blot analysis (separated on 12.5% polyacrylamide gels) of whole cells of *E. coli* strain DADE-P producing the indicated Cys substitutions in TatA and TatC (and wild type TatB, from plasmid pUNITATCC4) following exposure to 1.8mM CuP (oxidizing) or 10 mM DTT (reducing) for 1 min. Crosslinked products were visualized by immunoblotting using anti-TatC antibodies. The asterisk in H indicates a faint TatAC crosslink. **B.** The TatA^L9C^-TatC^F213C^ oxidized (O) and reduced (R) samples from A. were separately probed with an anti-TatA antibody (note that the TatA monomer that is in large excess has been run off the bottom of the gel). **C and F.** Cells of strain DADE harboring plasmid pTAT101 producing **C.** TatA^L9C^ and wild type TatB along with either TatC^V212C^ or TatC^F213C^, or **F.** TatA^S5C^, wild type TatB and TatC^F213C^, as indicated, were incubated with 1.8mM CuP (O) or 10 mM DTT (R) for 1 min. **D**. Structural model of TatA interacting with TatC at the TatA constitutive binding site. The backbone distances between TatA^S5/L9^/TatC^F213^, TatA^I6^/TatC^V212^, TatA^A13^/TatC^I220^ and TatA^V17^/TatC^E227^ are shown. **I.** Cells of strain DADE producing TatA^L9C^ and wild type TatB alongside TatC^F94A,E103A,M205C^ or TatC^F94A,E103A,F213C^ (annotated TatC^FEA,M205C^ or TatC^FEA,F213C^, respectively) from pTAT101 were left untreated (C) or incubated with 1.8mM CuP (O) for 1 min. For C. and D., following quenching, membranes were prepared, samples separated by SDS PAGE (12.5% polyacrylamide) and immunoblotted using an anti-TatC antibody.

Taken together, the absence of a TatA crosslink at the TatC polar cluster site, along with clear crosslinks between TatA^L9C/L11C^ and the N-terminal end of TatC TM6 suggests that TatA occupies a distinct binding site. Co-evolutionary analysis identified a weak evolutionary coupling between TatA/B residue 17 and TatC residue 227 (*E. coli* numbering), that was much lower than the primary contacts identified by Alcock *et al.* (2016). Guided by this coupling and the TatA/TatC crosslinks identified above we were able to dock TatA into a binding site that lies adjacent to the polar cluster site (Fig 3D; Fig 4). Atomistic molecular dynamic simulations suggested that TatA was stable in this site (Fig S4A) and together with the modelling predicted further contacts between TatA and TatC including S5-F213, I6-V212 and A13-I220. To confirm this we constructed cysteine substitutions at each of these predicted pairs, and were able to detect oxidant-induced TatA-TatC heterodimers at each of these positions (Fig 3E-G). We were also able to detect a very faint heterodimeric crosslink between TatA^V17C^ and TatC^E227C^ (Fig 3H). We conclude that TatA occupies a binding site that is distinct from, but adjacent to, the polar cluster site.

**Figure 4.**
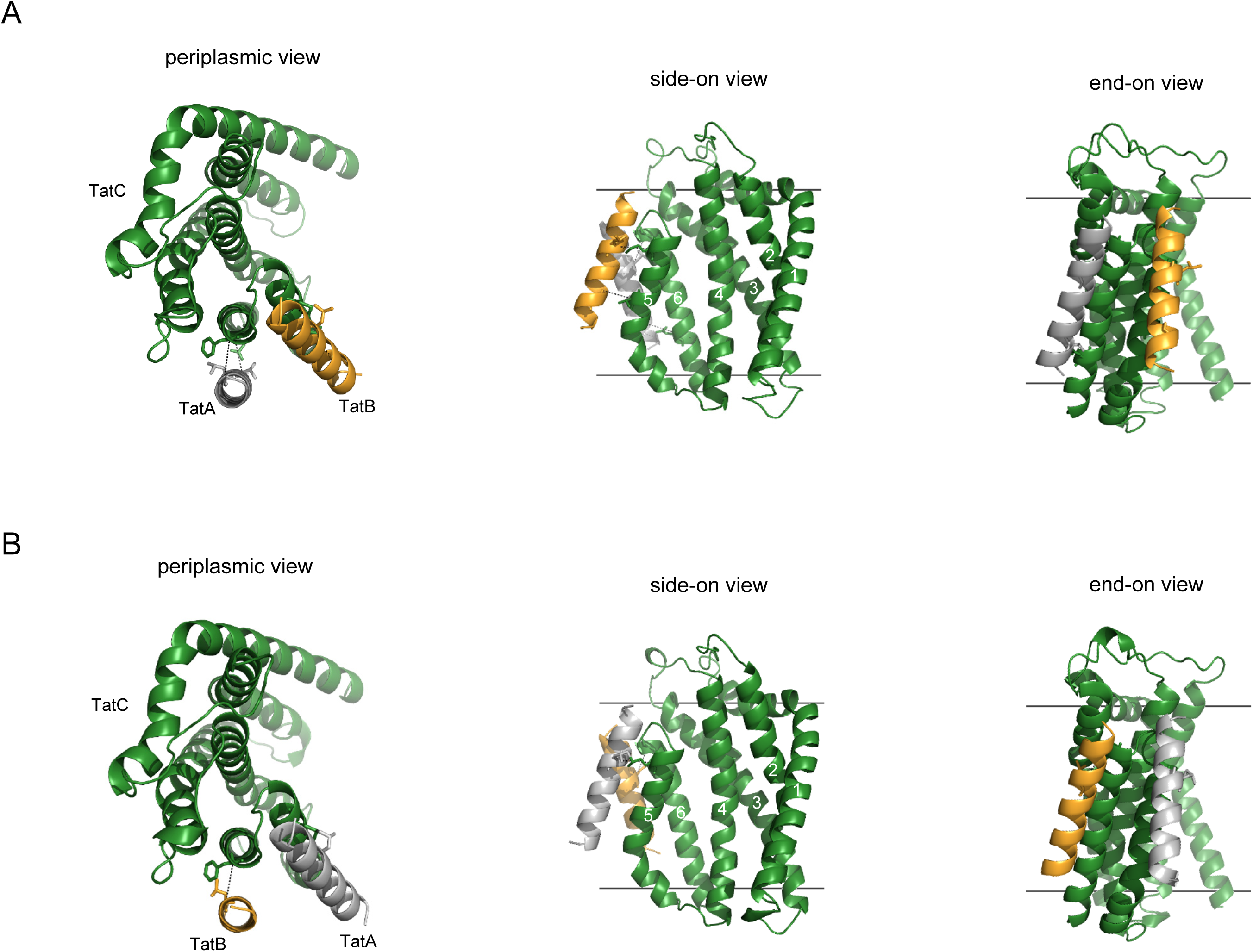
Models of the TatABC trimer in the resting and activated state. Three views of **A.**the resting state TatABC complex, and **B.** the substrate-activated TatABC complex. TatA is shown in silver, TatB gold and TatC green. Note that in B. the substrate signal peptide is not shown as it is currently unclear precisely where it binds in the activated state.

TatA associates with TatC in two different modes. One of these is constitutive, whereas the second is induced in the presence of substrate and is associated with Tat transport (Alcock et al., 2016; Aldridge et al., 2014; Blummel et al., 2015; Mori and Cline, 2002; Zoufaly et al., 2012). To determine whether TatA was constitutively bound at this newly-identified site, we introduced the F94A, E103A substitutions into TatC^F213C^ to prevent substrate binding and probed for interaction with TatA^L9C^. Fig 3I shows that the TatAC crosslink was still strongly detected, and therefore arises due to constitutively-bound TatA.

Collectively these results demonstrate that there are two binding sites for TatA family proteins on *E. coli* TatC and that under resting conditions TatB occupies the polar cluster site while TatA is bound at the TatA constitutive site. Molecular modelling indicates that both of these sites can be simultaneously occupied on a single TatC (Fig 4A). Atomistic molecular dynamics suggest that this ternary complex is stable as a TatA_1_B_1_C_1_ heterotrimer (Fig S4A), with stability further increased for an TatA_3_B_3_C_3_ oligomer (Fig S4B), and structural stability plots indicate that the secondary structure in the starting models was preserved (Fig S6A,C).

### TatA and TatB are each capable of occupying both binding sites

We next asked the question whether TatA and TatB were each capable of occupying these two binding sites if they were the only TatA-family protein present. Fig 5A shows that in the absence of TatB, a Cys substitution at TatA^L9^ still disulfide crosslinks with TatC^F213C^, indicating that it occupies its constitutive site. However, additional crosslinks were now also detected between TatA^L9C^ and TatC^M205C/L206C^ which are adjacent to the polar cluster site. This finding indicates that TatA is capable of binding in both sites. It is noteworthy in these experiments that the pattern of TatC homodimerization was also altered in the absence of TatB. Comparing Fig 3A with Fig 5A, a strong TatC^M205C^ homodimer was detected when TatA rather than TatB was present at the polar cluster site. Likewise, the pattern of homodimeric crosslinks seen in the TatC P3 loop differed in the presence and absence of TatB (for example TatC^T208C^ gave a stronger self-crosslink than TatC^P209C^ or TatC^P210C^ in the presence of TatB, whereas the reciprocal was observed when TatB is absent).

**Figure 5.**
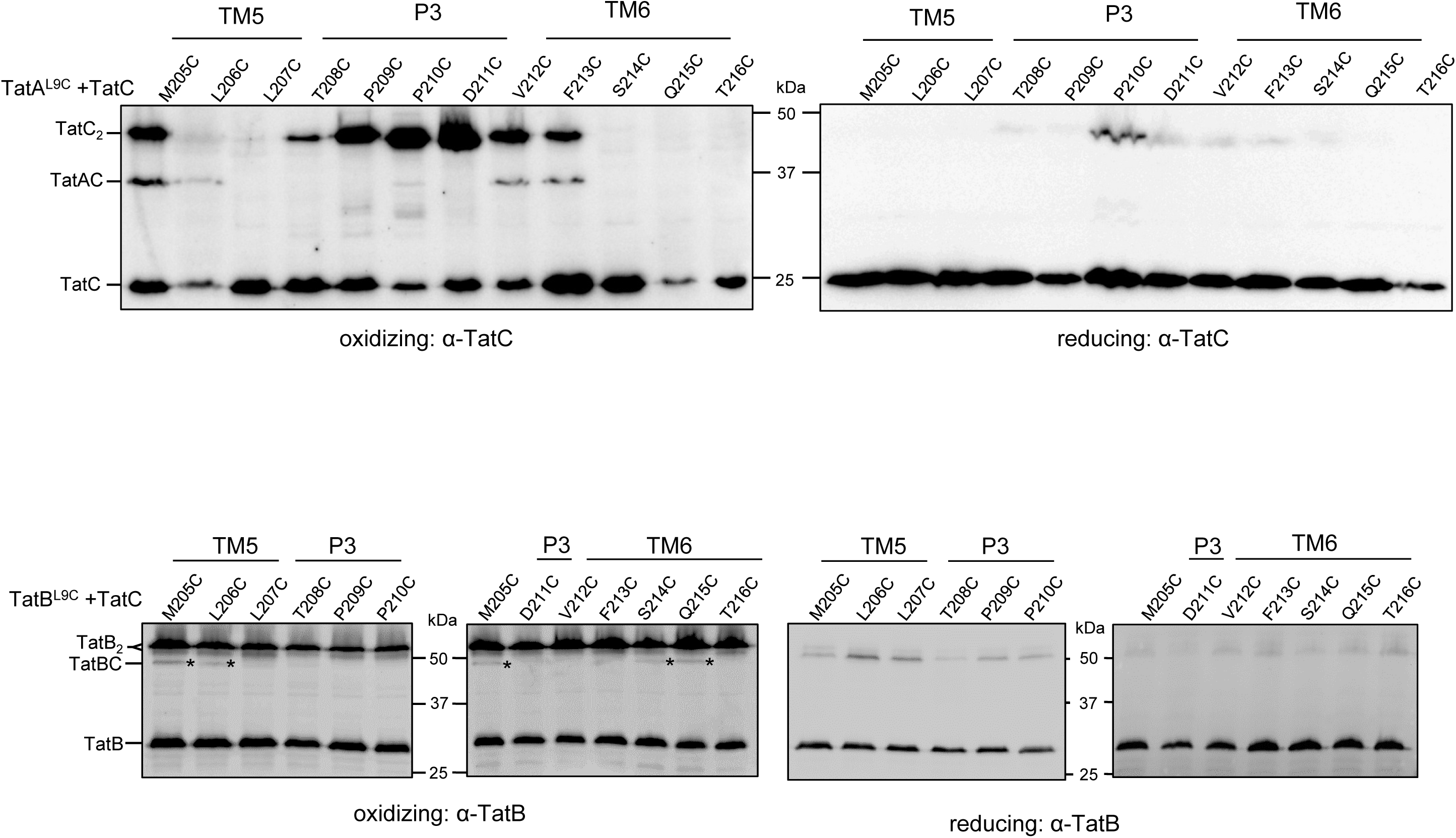
TatA and TatB can each occupy both binding sites on TatC. A.Western blot analysis (separated on 12.5% polyacrylamide gels) of whole cells of *E. coli* strain DADE-P producing TatA^L9C^ alongside the indicated Cys substitutions in TatC (in the absence of TatB, from plasmid pUNITATCC4∆B) following exposure to 1.8mM CuP (oxidizing) or 10 mM DTT (reducing) for 1 min. Crosslinked products were visualized by immunoblotting using anti-TatC antibodies. B. Western blot analysis (separated on 10% polyacrylamide gels) of membranes from *E. coli* strain DADE producing TatB^L9C^ alongside the indicated Cys substitutions in TatC (from plasmid p101C*BC) following exposure of whole cells to 1.8mM CuP (oxidizing) or 10 mM DTT (reducing) for 1 min. Crosslinked products were visualized by immunoblotting using an anti-TatB peptide antibody. The asterisks indicate likely TatBC crosslinks.

When TatA was absent, crosslinks of TatB^L9C^ to TatC^M205C^ and TatC^L206C^ were still detected, indicating occupancy at the polar cluster site, but additional crosslinks were also now detected between TatB^L9C^ and TatC^S214C/Q215C^, showing that TatB can also bind close to the TatA constitutive site if this site is vacant. Thus we conclude that each protein is able to occupy both binding sites.

### TatA and TatB switch binding sites in the presence of a Tat substrate

We next addressed whether differential occupancy of TatA and TatB at these binding sites was functionally related to Tat transport. To this end we undertook disulfide crosslinking analysis in the presence of an overproduced Tat substrate, CueO. We focused initially on the interaction of TatB^L9C^ and TatC^M205C^ that reports on the presence of TatB at the polar cluster site. When CueO was overproduced, the level of crosslinking between TatB^L9C^ and TatC^M205C^ appeared to diminish, and the level of TatB and TatC homodimers to increase compared to those seen in the presence of endogenous substrate proteins (Fig 6A). It should be noted, in agreement with this, that substrate-induced TatC homodimerization through M205C has previously been observed (Cleon et al., 2015). This finding is consistent with the idea that there is substrate-induced movement of TatB^L9^ away from TatC^M205^. To determine whether TatB may now occupy the second TatA/TatB binding site, we probed for crosslinks between TatB^L6C^, TatB^L9C^, TatB^L10C^ or TatB^L11C^ and Cys substitutions at positions 213, 216 or 217 of TatC, each in the presence of overproduced CueO. Fig 6B shows that a crosslink could be detected between TatB^L9C^ and TatC^F213C^. It was noted in Fig 2 that in the presence of endogenous substrate a very faint crosslink between these same two residues could be detected with the anti-TatB antibody (but not the anti-TatC antibody). The observation that this crosslink was abolished when substrate binding to the TatBC complex was prevented by introduction of the TatC^F94A,E103A^ substitutions (Fig 2D), even when substrate was overproduced (Fig 6C) strongly suggests that docking of a substrate to the receptor complex triggers the movement of TatB into the TatA constitutive binding site.

**Figure 6.**
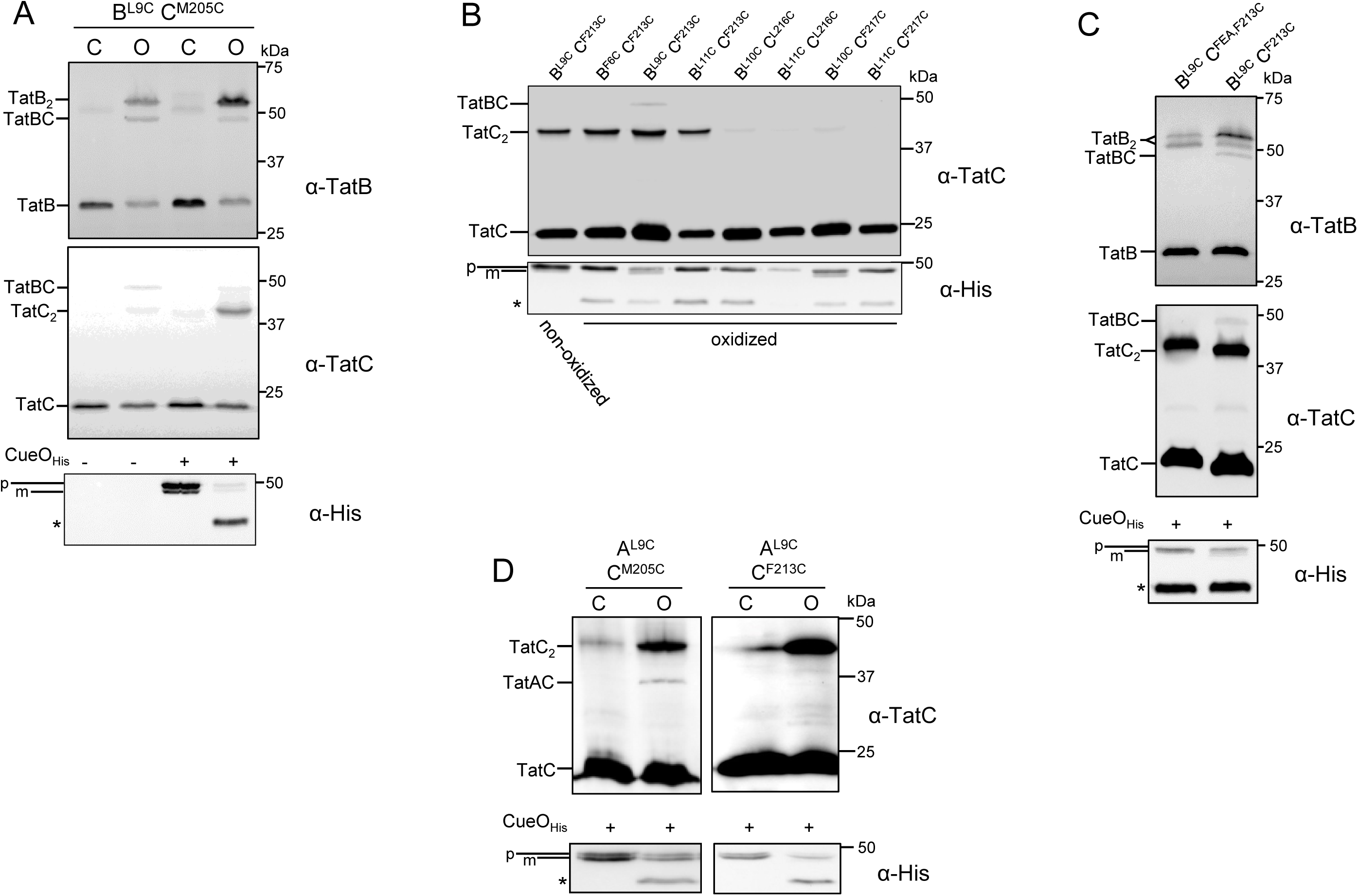
TatA and TatB crosslinking patterns are altered in the presence of an overproduced Tat substrate. A. Strain MC4100∆BC harboring plasmid p101C*BC producing TatB^L9C^ alongside TatC^M205C^ and plasmid pQE80-CueO where indicated, were left untreated (Control, C), or incubated with 1.8mM CuP for 1 min (O). Membrane fractions were prepared, separated by SDS PAGE (10% polyacrylamide) and immunoblotted with anti-TatB_FL_, anti-TatC as indicated. An aliquot of the soluble fraction following membrane preparation was retained and analyzed by immunoblotting with an anti-Histag antibody. **B.** Strain MC4100∆BC producing the indicated Cys variants of TatB and TatC from plasmid p101C*BC and his-tagged CueO from pQE80-CueO were incubated with 1.8mM CuP for 1 min. Following quenching membrane fractions were separated by SDS PAGE (10% polyacrylamide) and immunoblotted with an anti-TatC antibody. A non-oxidized sample of membranes harboring TatB^L9C^ - TatC^MF213C^ is shown in the left-most lane. An aliquot of the soluble fraction from each sample was retained and analyzed by immunoblotting with an anti-Histag antibody. **C.** Whole cells of strain MC4100∆BC producing TatB^L9C^ alongside TatC^F213C^ or TatC^F94A,E103A,M205C^ (annotated TatC^FEA,M205C^) from plasmid p101C*BC, and histagged CueO (from pQE80-CueO) were incubated for 1 min with 1.8mM CuP. Following membrane preparation, crosslinks were detected with an anti-TatB peptide antibody or an anti-TatC antibody. An aliquot of the soluble fraction from each sample was retained and analyzed by immunoblotting with an anti-Histag antibody. **D.** Strain DADE harboring plasmid pTAT101 producing wild type TatB, TatBA^9C^ and either TatC^M205C^ or TatC^F213C^ along with plasmid pQE80-CueO were left untreated (Control, C), or incubated with 1.8mM CuP for 1 min (O). Membrane fractions were separated by SDS PAGE (12.5% polyacrylamide) and immunoblotted with an anti-TatC antibody. An aliquot of the soluble fraction from each sample was retained and analyzed by immunoblotting with an anti-Histag antibody. p: precursor, m: mature forms of substrate CueO-His * indicates an oxidation product of CueO.

Since TatB can occupy the TatA constitutive site in the presence of overproduced substrate, it should be accompanied by loss of TatA from this binding site. Fig 6D shows, in agreement with this, a crosslink between TatA^L9C^ and TatC^F213C^ could no longer be detected when CueO was overproduced. Instead a distinct crosslink could now be seen between TatA^L9C^ and TatC^M205C^ indicating that TatA has moved to occupy the polar cluster binding site upon substrate binding. Thus substrate binding appears to trigger TatA and TatB to switch positions.

Molecular dynamic simulations have previously shown that TatA can stably interact with TatC through the polar cluster site (Alcock et al., 2016). A similar analysis indicates that TatB can bind at the TatA constitutive site and that it can stably occupy that site when TatA is bound at the polar cluster site (Fig S5, S6B). A model for the substrate-activated state of the TatABC complex is shown in Fig 4B.

## Discussion

In this study we have used disulfide crosslinking to probe the interaction of TatA and TatB with TatC under resting conditions and in the presence of an overexpressed substrate. Our results have delineated two binding sites for these proteins. One of these – the ‘polar cluster’ site - has been identified previously and involves key interactions between a polar side chain at position 8 of TatA/B and a patch of conserved residues M205, T208 and Q215 in TatC (Alcock et al., 2016). The second site lies adjacent to the polar cluster site, at TM6 of TatC. Experiments where TatA or TatB were present individually as the sole TatA/B family protein indicated that each of these proteins was capable of occupying both sites. However, locking the Tat system into the resting state through the introduction of substitutions in TatC that prevent signal peptide binding demonstrated that TatB occupies the polar cluster site under these circumstances, in agreement with the conclusions of Alcock *et al.* (2016), with TatA occupying the newly-identified site. Modelling and molecular dynamics simulations showed that interaction of TatA with its constitutive binding site was stable and that both of these sites can be simultaneously occupied on one TatC protein. This adjacent positioning of TatA and TatB is supported by the detection of TatA-TatB crosslinks when a photocrosslinker is introduced into the N-terminal region of TatB (Blummel et al., 2015).

Recently, a molecular model of the multivalent resting-state TatBC complex was built by docking TatBC protomers together using evolutionary couplings between TatC proteins and crosslinks between the TatC TM1 and the TatB TM (Alcock et al., 2016; Blummel et al., 2015). Fig 7 shows updated models for the resting TatABC complex containing either three or four copies of the heterotrimer. TatA can be readily accommodated into the complex with minimal adjustment, slotting into a groove that is present at the outside of the complex. This peripheral binding of TatA likely explains why a TatBC complex can be stably purified when TatA is absent (e.g. (Behrendt et al., 2007; Orriss et al., 2007) and may potentially account for findings that TatA is variably shed from the TatBC complex during purification in detergent solution (Bolhuis et al., 2001; McDevitt et al., 2006).

**Figure 7.**
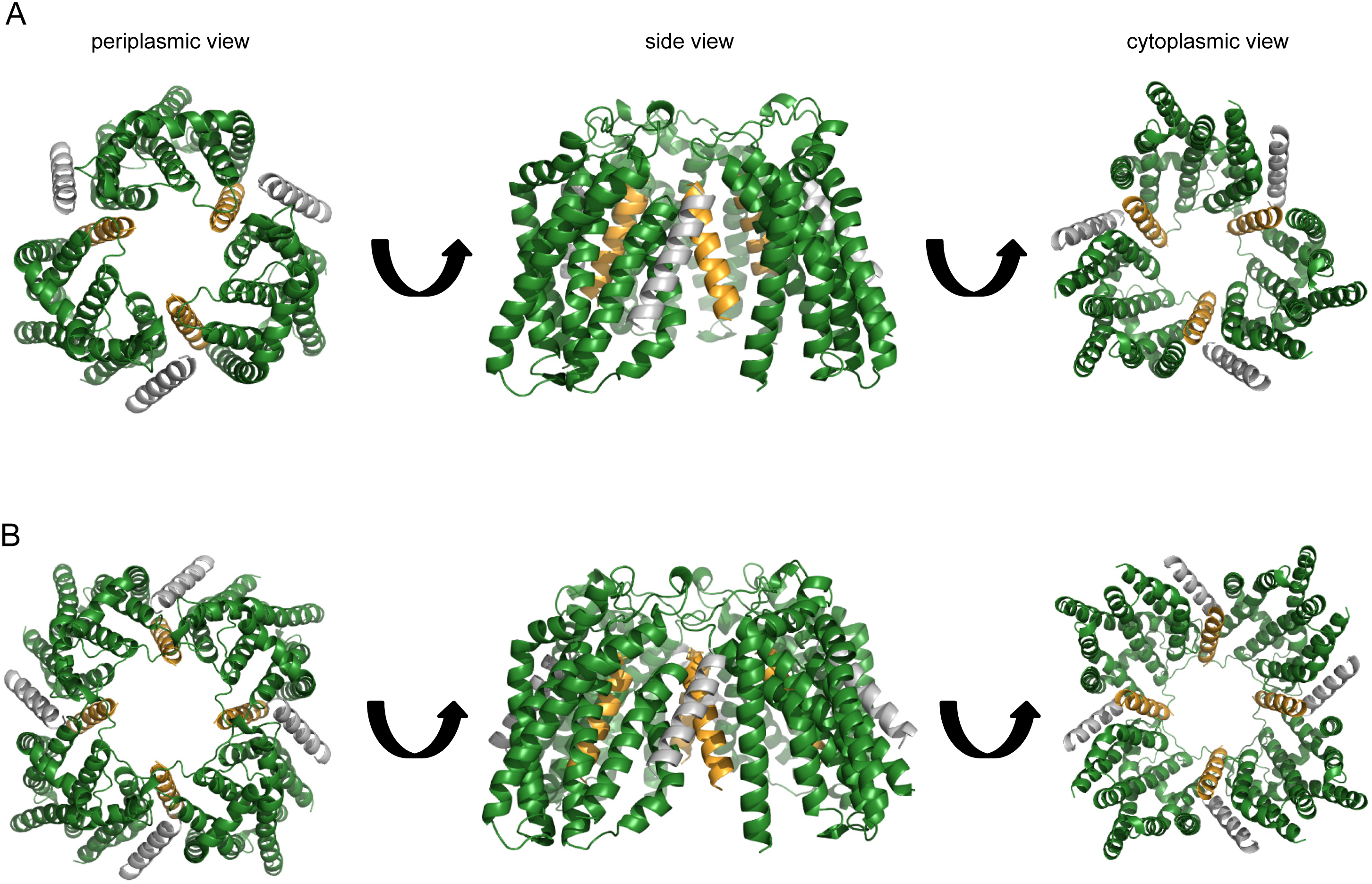
Models of the multimeric resting state TatABC complex. Models based on A. three or B. four heterotrimers. Modified from (Alcock et al., 2016).

When a Tat substrate protein is overproduced, we show that TatA can no longer be crosslinked in the constitutive binding site but is instead detected at the polar cluster site. This is accompanied by a reduction in the level of TatB crosslinking at the polar cluster site and the appearance of TatB-TatC crosslinks at the TatA constitutive site. In agreement with this, substrate-dependent contacts between the chloroplast TatA ortholog, Tha4 and cpTatC at the equivalent polar cluster site have also been observed (Aldridge et al., 2014). Position-switching of TatA and TatB is likely triggered by signal peptide binding at the complex. In the deep-binding mode, contacts have been detected between the signal peptide h-region and both the TatB TM and TatC TM5, close to the polar cluster site (Alami et al., 2003; Aldridge et al., 2014; Blummel et al., 2015). While current findings cannot distinguish whether it is TatA or TatB that makes the initial movement, we speculate it is TatB. It has been shown that Tat signal peptides are sequestered within a cavity comprising TatB and TatC, which could potentially correspond to the central cavity seen in the modelled TatBC/TatABC complexes (Fig 7 and (Alcock et al., 2016; Blummel et al., 2015). Note that the polar cluster site is adjacent to the lumen of this cavity and, accordingly, docking of the signal peptide close to this region may cause conformational rearrangements that drive TatB from the polar cluster site into the TatA constitutive site. Further support for this suggestion comes from a recent genetic study where suppressors of inactive twin arginine signal peptides or a defective signal peptide binding site were identified that located to the TM of TatB. Crosslinking analysis indicated that at least some of these TatB variants caused rearrangement at the polar cluster site, probably by decreasing TatB binding affinity at this site and/or increasing binding affinity for the constitutive site (Huang et al., 2017).

In addition to changes in TatA and TatB crosslinking patterns, we note that signal peptide binding also resulted in the formation of TatB homodimers through L9C, and TatC homodimers mediated through TatC M205C or through F213C. The formation of a substrate-induced TatC M205C homodimer has been observed previously and taken to report on the activated state of the Tat translocase (Cleon et al., 2015)(Huang et al 2017). The head-to-tail arrangement of TatC in the resting state model of TatABC complex positions neighboring TatC M205 residues 25Å away from each other, and F213 residues even further apart, a distance that is too great for disulfide bond formation through Cys sidechains at these positions. This strongly suggests that there must be a significant conformational change in the TatABC complex upon substrate binding to bring TatC protomers into a tail-to-tail organisation. Opening up of the complex in this way would then allow TatA to access the vacated polar cluster site. It should be noted that the concave face of TatC has been implicated in the nucleation of the transport-active TatA oligomer (Aldridge et al., 2014; Rollauer et al., 2012). Binding of a TatA molecule at the polar cluster site places it adjacent to the concave face where it could potentially initiate polymerization of further TatA molecules.

In conclusion we have defined two binding sites for TatA family proteins within the TatABC complex and have demonstrated differential occupancy of TatA and TatB at these sites during different stages of Tat transport. These findings help to explain a long-standing observation that overproduction of TatB relative to TatA and TatC inactivates the Tat system (Sargent et al., 1999) since simultaneous occupancy of TatB (which is normally present 20 fold less than TatA; Jack et al., 2001; Sargent et al., 2001) in both binding sites would be expected to block progression through the transport cycle.

## Materials and Methods

### Strains and plasmids and growth conditions

All strains used for crosslinking analysis are derived from MC4100 (F^−^, *[araD139]_B/r_, Δ*(*argF-lac*)*U169, λ^-^, e14-, flhD5301, ∆*(*fruK-yeiR*)*725*(*fruA25*), *relA1, rpsL150*(Str^R^*), rbsR22, ∆*(*fimB-fimE*)*632*(*::IS1*), *deoC1* - (Casadaban and Cohen, 1979). MC4100∆BC (as MC4100, *∆tatBC* (Alcock et al., 2013)), DADE (as MC4100, *∆tatABCD ∆tatE*); (Wexler et al., 2000) and DADE-P (as DADE, *pcnB1 zad-*981::Tn*10*d (Kan^r^); (Lee et al., 2006)) were used where indicated in the figure legends. Strain JM109 (F' *traD36 proA^+^B^+^ lacI^q^ ∆*(*lacZ*)*M15/∆*(*lac-proABI glnV44 e14^-^ gyrA96 recA1 relA1 endA1 thi hsdR17)* was used for cloning purposes.

All plasmids used in this study are listed in Table S1. Plasmid pUNITATCC4 encodes TatA, TatB and cysteine-less TatC in plasmid pQE60. Production of the encoded proteins is driven by the phage *T5* promoter which is constitutively active in strains deleted for *lacI,* such as MC4100 derivatives. Plasmid pUNITATCC4∆B (producing TatA and TatC) was derived from pQEA(DB)C (Fritsch et al., 2012) by excising DNA covering the wild type allele of *tatC* through digestion with *Xho*I and *Bam*HI and replacement with a Cys-less *tatC* allele amplified using oligonucleotides TatBdeldownXho (Fritsch et al., 2012)and TatCBam (Lee et al., 2006) with pUNITATCC4 as template. Plasmid pTAT101 codes for TatA, TatB and TatC on a low copy number vector and produces these proteins at approximately four times chromosomal level (Kneuper et al., 2012). Plasmid p101C*BC expresses *tatBC* at approximately chromosomal level and has been described previously (Alcock et al., 2013). p101C*BC Cys-less was designed as follows: a *tatBC* allele where all of four Cys codons of *tatC* had been mutated to Ala codons was amplified from pTat101 Cys less (Cleon et al., 2015) using primers BamHI-TatB-F and SpHI-TatC-R (Table S2), introducing a *Sph*I site at the 3’-end of *tatC.* The PCR product was digested with *Bam*HI*/Sph*I and cloned into similarly-digested p101C*BC. All point mutations in plasmids were introduced by Quickchange site-directed mutagenesis (Stratagene) using the primers listed in Table S2. Plasmid pQE80-CueO expresses *E. coli* CueO with a C-terminal his6 tag and has been described previously (Leake et al., 2008).

Phenotypic growth in the presence of 2% SDS was assessed by culturing strains of interest in LB medium containing appropriate antibiotics until an OD_600_ of 1 was reached, after which 5μL aliquots of culture were spotted onto agar plates containing LB or LB supplemented with 2% SDS and appropriate antibiotics. Plates were incubated at 37°C for 16 hours after which they were photographed. Antibiotics were used at the following concentrations: chloramphenicol (25 μg.ml^−1^), kanamycin (50 μg.ml^−1^) and ampicillin (125 μg.ml^−1^).

### *In vivo* disulfide cross-linking experiments

For Tat proteins produced at close to native level (from pTAT101 and p101C*BC), the appropriate *E. coli* strain/plasmid combination was cultured overnight in LB medium containing appropriate antibiotics. Cells were diluted 1:100 into fresh LB medium supplemented with appropriate antibiotics and cultured aerobically until an OD_600_ of 0.3 was reached. For the CuP titration experiment, six 25 mL aliquots were withdrawn and each supplemented with fresh LB medium to a final OD_600_ of 0.15. The first aliquot was left untreated (control), the second one was supplemented with 10 mM DTT (reducing) and the remainder were incubated with 0.3, 0.6, 1.2 or 1.8 mM CuP (oxidising). Cells were incubated for 15 min at 37°C with agitation, then harvested, resuspended in 1 mL 20 mM Tris-HCl, pH 7.5, 200 mM NaCl, 12 mM EDTA, 8 mM N-ethylmalemide and incubated at 37°C for 10 minutes to quench free sulphydryls. The cell suspension was supplemented with protease inhibitor cocktail (Roche) and disrupted by sonication. Unbroken cells were removed by centrifugation (10 000 × *g* for 5 min at 4°C) and the supernatant ultracentrifuged (200 000 × *g* for 30 min at 4°C). The membrane pellet was resuspended in 70 μL 1 × Laemmli buffer lacking β-mercaptoethanol (BioRad). For the time course experiment, when subcultured cells reached OD_600_ of 0.3, seven 25 mL aliquots were withdrawn and supplemented with fresh LB medium to a final OD_600_ of 0.15. One aliquot was left untreated, one was supplemented with 10 mM DTT and the remainder incubated with 1.8mM CuP for 1, 2, 5, 10 or 15 min at 37°C with agitation. The reactions were quenched for 10 min as before and a small aliquot of cells from each sample was withdrawn, serially diluted and spread on LB plates supplemented with appropriate antibiotics to assess viability. Membrane samples were prepared from the remainder of the cells and treated as described above. For all other experiments, when cells reached OD_600_ 0.3, three separate 25 mL aliquots were withdrawn and supplemented with fresh medium to OD_600_ of 0.15. One aliquot was left untreated, the second supplemented with 10 mM DTT and the third incubated with 1.8mM CuP for 1 min at 37°C with agitation. The reactions were quenched and membrane samples prepared from the as described before. When experiments were performed in the presence of overproduced CueO, cells additionally harboured pQE80-CueO and 1 mM IPTG was included in the initial subculture.

For Tat proteins produced at higher copy from plasmid pUNITATCC4, overnight cultures of DADE-P harbouring pUNITATCC4 were subcultured at 1:100 to inoculate fresh LB containing appropriate antibiotics. When cells reached OD_600_ of 0.3, three 2.5 mL aliquots were withdrawn and made up to 5 mL with fresh LB to a final OD_600_ of 0.15. These aliquots were treated and quenched as described above after which the cells were harvested at 16000 × *g* for 1 min and resuspended in 40 μL of 1 × Laemmli buffer lacking β-mercaptoethanol (BioRad).

For analysis, sodium dodecyl sulfate polyacrylamide gel electrophoresis (SDS-PAGE) was performed using Tris-glycine gels (Laemmli, 1970). 20 μl of sample was analyzed in each case. Following electrophoresis, proteins were transferred to nitrocellulose membrane (I-blot^®^ system, Life Technologies). TatA, and TatC were identified using the polyclonal antibodies previously described (Cleon et al., 2015; Sargent et al., 2001). Two different TatB antisera were used. One of these was raised against full length TatB (Sargent et al., 2001) and in this study is annotated as TatB_FL_. The second was raised against two peptides of *E. coli* TatB 69-84 and TatB 156-171, and was then affinity purified with peptide 156-171 (Alcock et al., 2016) and in this study is referred to as a TatB peptide antibody. Abcam Anti-6X His tag^®^ antibody [GT359] (HRP conjugate) was purchased (catalogue number ab184607), and a HRP-conjugated goat anti-rabbit antibody (BioRad, catalog number 170-6515) was used as secondary antibody for the TatA, TatB and TatC antisera. Cross-reacting bands were visualised after incubation with Clarity™ Western ECL Blotting Substrate (Biorad) using a CCCD camera (GeneGnome XRQ, Syngene).

### Molecular modelling and simulations

Molecular modelling was carried out as described previously (Alcock et al., 2016). All images were generated using Pymol (The PyMol Molecular Graphics System, Version 1.8, Schrödinger, LLC). Multimers were built using TatA-TatC/TatB-TatC disulfide crosslinks as unambiguous constraints for docking using Haddock (Dominguez et al., 2003).

All MD simulations were performed using GROMACS v5.1.2 (Pronk et al., 2013). The Martini 2.2 force field (de Jong et al., 2013) was used to run initial 1 μs Coarse Grained (CG) MD simulations to permit the assembly and equilibration of 1-palmitoly, 2-oleoyl phosphatidylglycerol (POPG): 1-palmitoly, 2-oleoyl phosphatidylethanolamine (POPE) bilayers around the TatABC complexes at a 1:3 molar ratio (Stansfeld et al., 2015). CG molecular systems were converted to atomistic detail using CG2AT (Stansfeld and Sansom, 2011), with Alchembed used to remove any unfavourable steric contacts between protein and lipid (Jefferys et al., 2015). The heterotrimeric atomistic systems equate to a total size of ~80,000 atoms and box dimensions in the region of 125 × 125 × 100 Å^3^, while the heterononameric systems comprised ~115,000 atoms, with box dimensions in the region of 100 × 100 × 100 Å^3^. The systems were equilibrated for 1 ns with the protein restrained before 3 repeats of 100 ns of unrestrained atomistic MDS, for each configuration of the molecular system (see below), using the Gromos53a6 force field (Oostenbrink et al., 2004). Molecular systems were neutralised with a 150 mM concentration of NaCl.

All simulations were executed at 37 °C, with protein, lipids and solvent separately coupled to an external bath, using the velocity-rescale thermostat (Bussi et al., 2007). Pressure was maintained at 1 bar, with a semi-isotropic compressibility of 4 × 10^−5^ using the Parrinello-Rahman barostat (Parrinello and Rahman). All bonds were constrained with the LINCS algorithm (Hess et al., 1997). Electrostatics was measured using the Particle Mesh Ewald (PME) method (Darden et al., 1993), while a cut-off was used for Lennard-Jones parameters, with a Verlet cut-off scheme to permit GPU calculation of non-bonded contacts. Simulations were performed with an integration time step of 2 fs. Analysis was performed using Gromacs tools and locally written python and perl scripts.

## Supplemental Information

**Figure S1.**
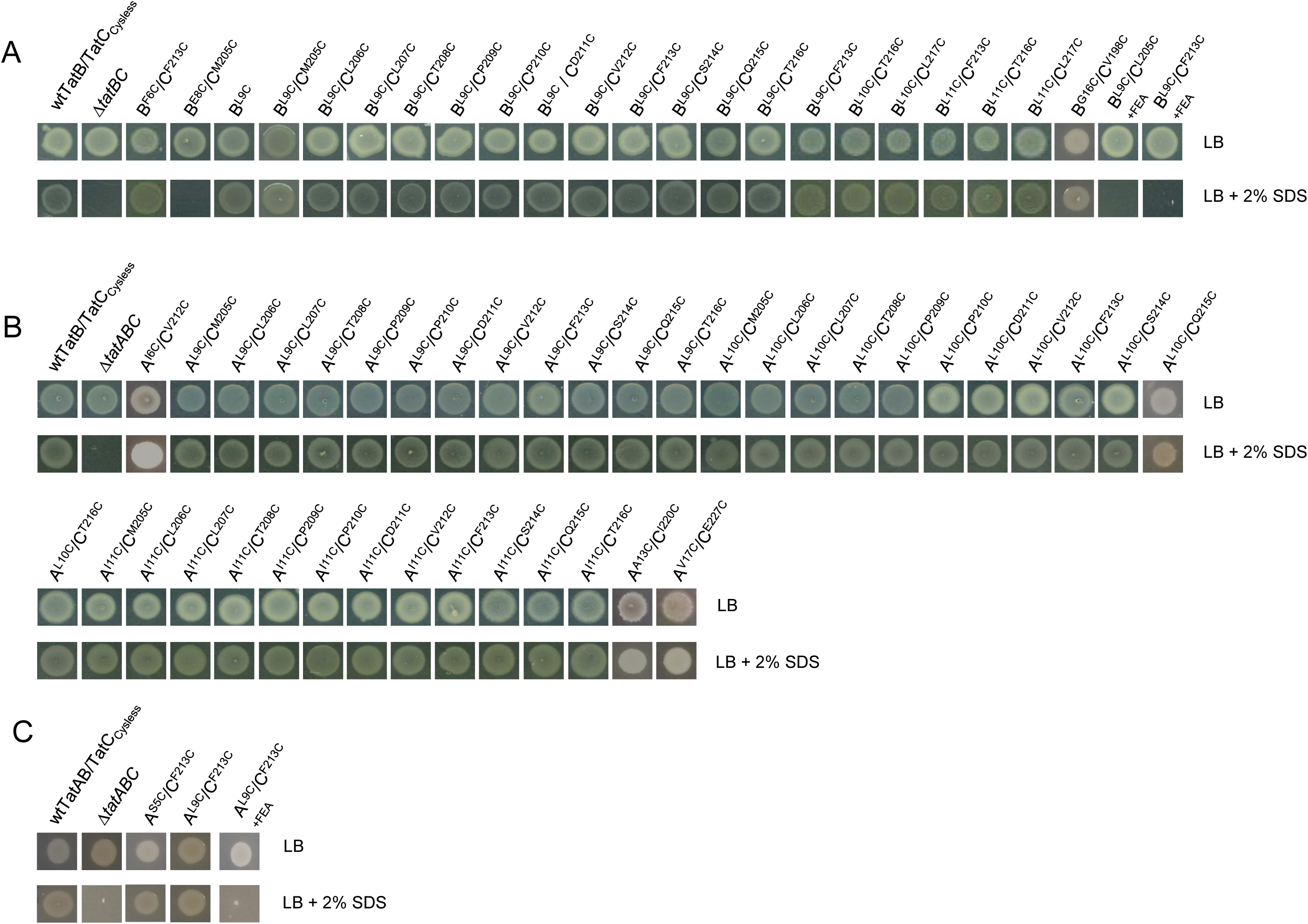
Phenotypic analysis of Tat activity in cells harbouring cysteine substitutions of TatA, TatB and TatC. A. TatB/TatC mutant pairs. Strain MC4100∆BC harboring either an empty vector (***∆****tatBC*) or plasmid p101C*BC encoding the indicated TatB and TatC variants alongside wild type TatA was spotted on LB medium or LB medium containing 2% SDS. **B.** and **C.** TatA/TatC mutant pairs **B.** Strain DADE-P harboring either an empty vector (***∆****tatABC*) or plasmid pUNITATCC4 encoding the indicated TatA and TatC variants alongside wild type TatB was spotted on LB medium or LB medium containing 2% SDS. **C**. Strain DADE harboring either an empty vector (***∆****tatABC*) or plasmid pTAT101 encoding the indicated TatA and TatC variants alongside wild type TatB was spotted on LB medium or LB medium containing 2% SDS. In each case An 8μl aliquot of each strain/plasmid combination following aerobic growth to an OD_600_ of 1.0 was spotted and incubated for 16 hr at 37°C prior to photographing.

**Figure S2.**
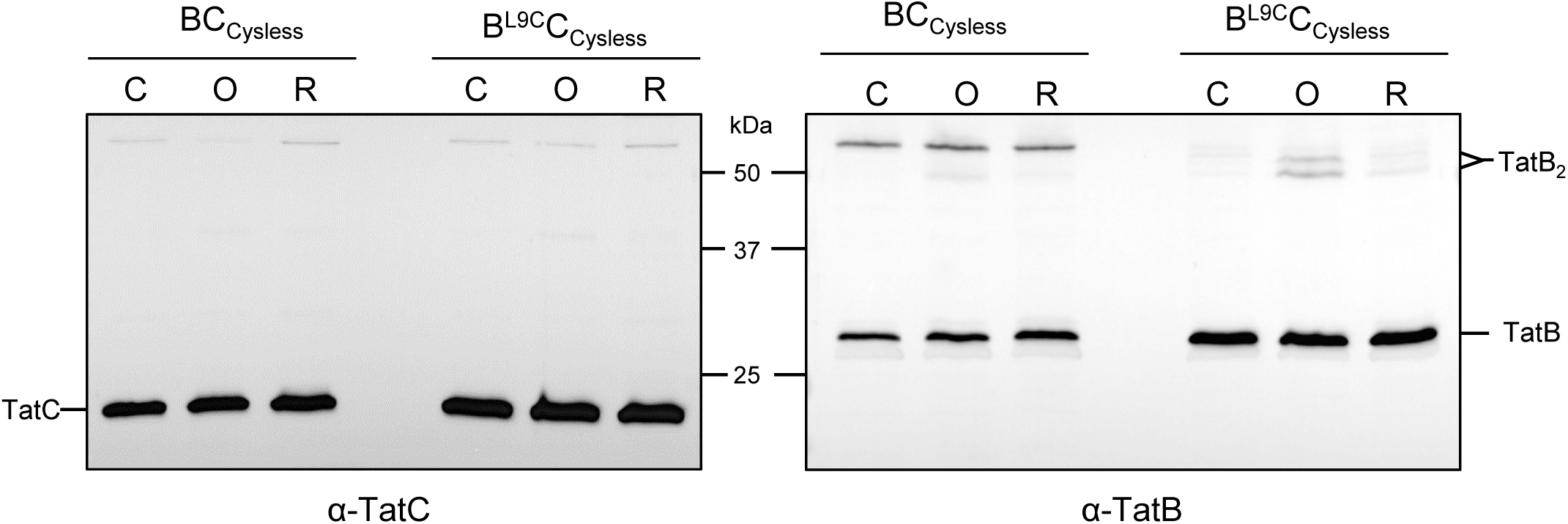
The TatB homodimer is recognized as a doublet band by the TatB antipeptide antibody. Western blot analysis of membranes from *E. coli* strain MC4100∆BC producing Cys-less TatC along with either native (Cys-less) TatB or TatBL9C, as indicated, from plasmid p101C*BC. Whole cells were either exposed to 1.8mM CuP (oxidizing; O) or 10 mM DTT (reducing; R) for 1 min, or left untreated (control; C). TatC and TatB were visualized by immunoblotting using anti-TatC or anti-TatB peptide antibodies.

**Figure S3.**
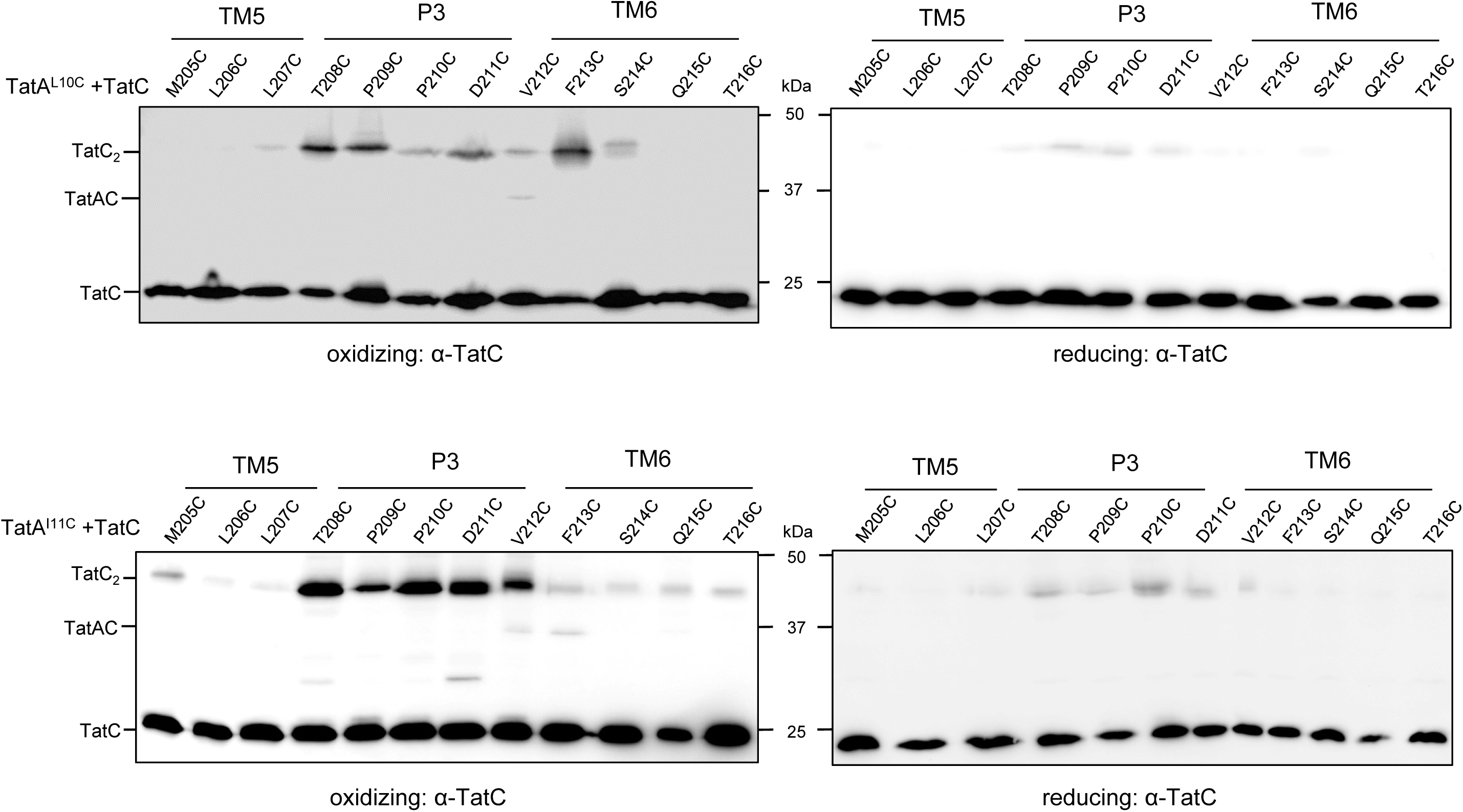
TatA^L10C^ interacts with TatC^v212C^ and TatA^I11C^ interacts with TatC^V212C/F213C^ *in vivo*. **A.** and **B.** Western blot analysis (separated on 12.5% polyacrylamide gels) of whole cells of *E. coli* strain DADE-P producing **A.** TatA^L10C^ or **B.** TatA^I11C^ alongside the indicated Cys substitutions in TatC (and wild type TatB, from plasmid pUNITATCC4) following exposure to 1.8mM CuP (oxidizing) or 10 mM DTT (reducing) for 1 min. Crosslinked products were visualized by immunoblotting using anti-TatC antibodies.

**Figure S4.**
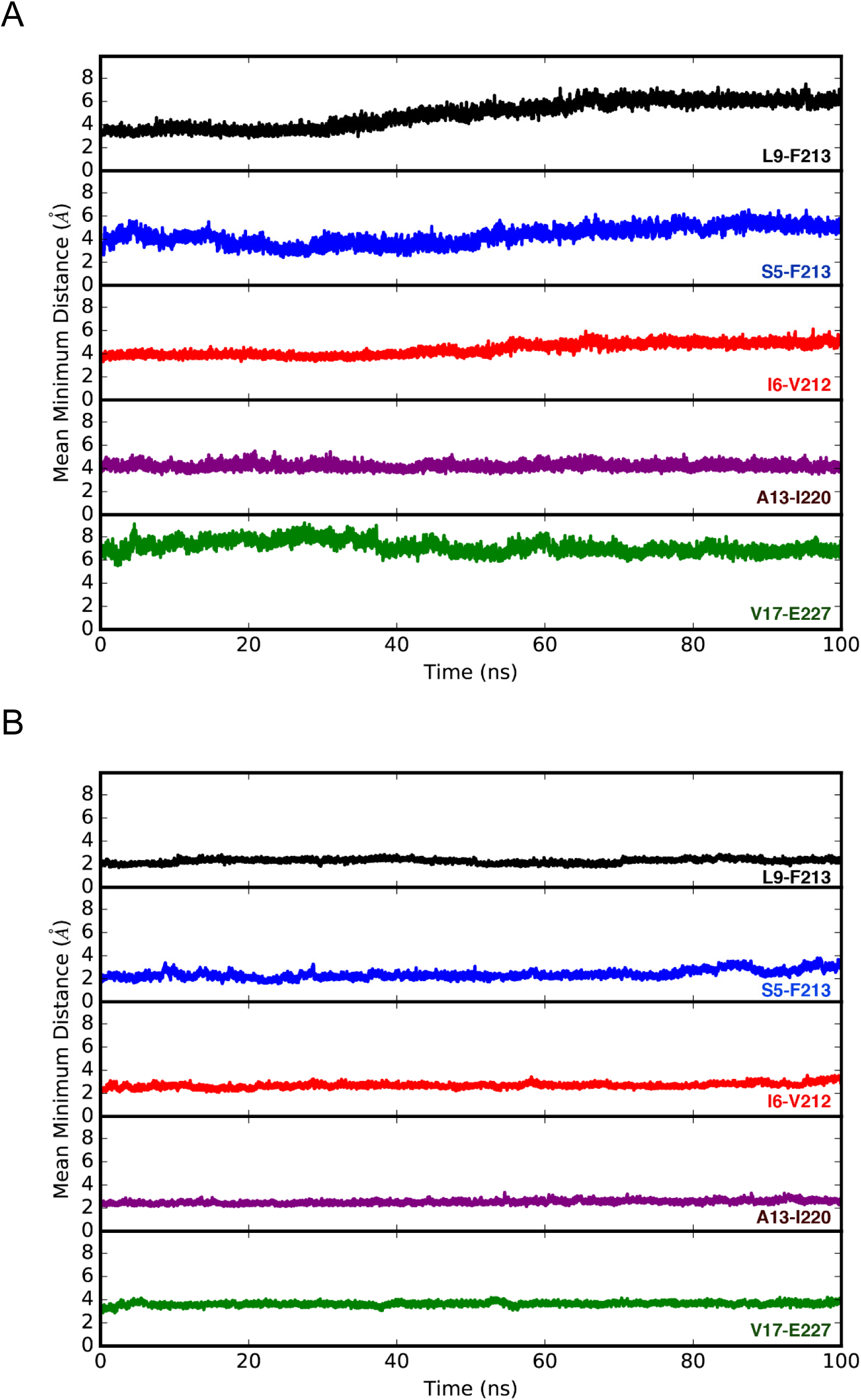
Molecular simulations of the interactions of TatA with TatC at its constitutive site and TatB with the TatC polar cluster site. TatA, TatB and TatC, assembled with TatB in the polar cluster site and TatA in its constitutive site, simulated for 100 ns as either **A.** Heterotrimeric TatA_1_B_1_C_1_ or **B.** Heterononameric TatA_3_B_3_C_3_. The plots show the mean minimum distance over time between the residues of TatA and TatC that are shown to be able to form cross-links. In both instances the residues remain in close proximity to one another, however, the TatA_3_B_3_C_3_ complex appears to stabilise the overall motions of the individual subunits and therefore the distances between residues remain consistently low.

**Figure S5.**
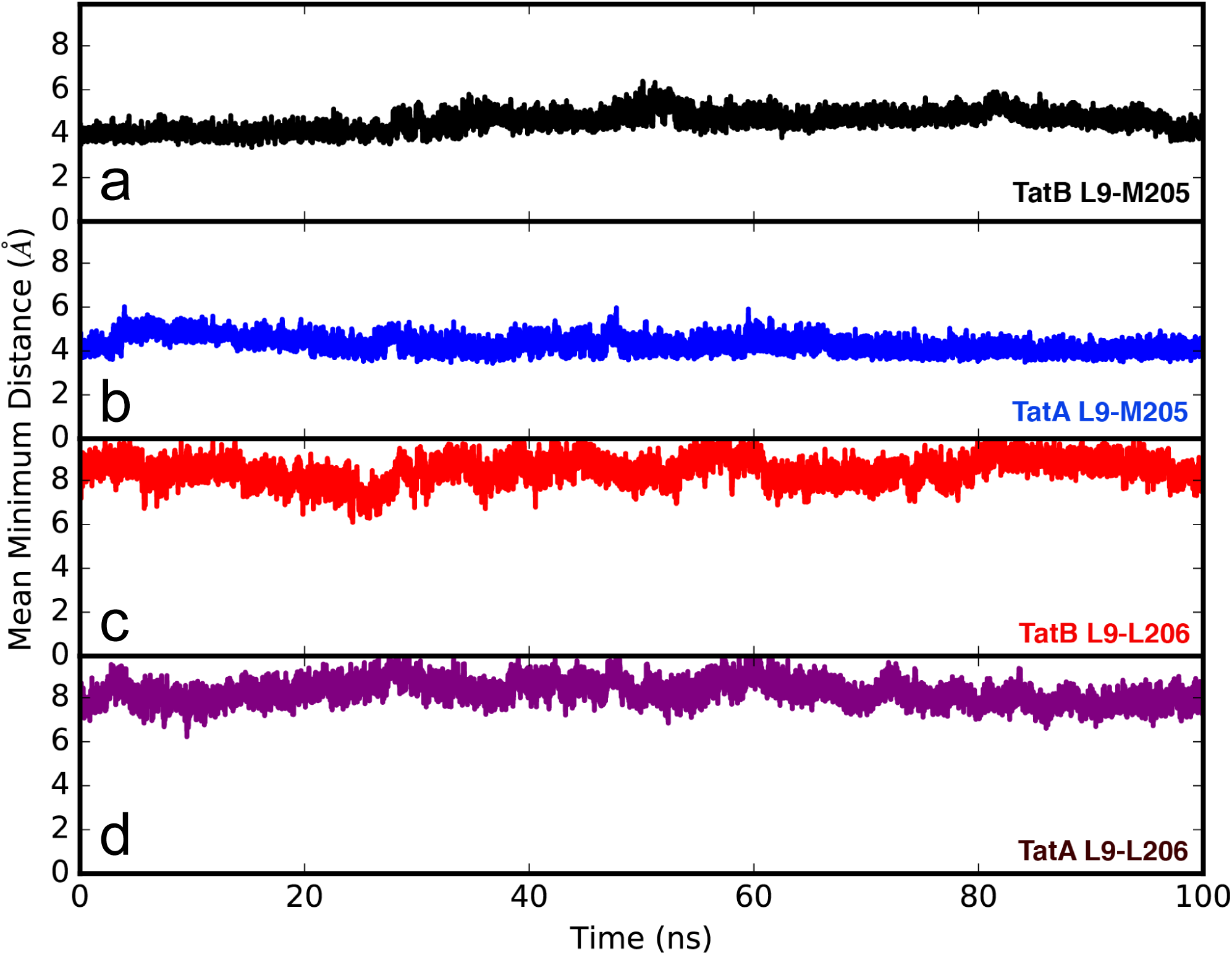
Molecular simulations of the interactions of TatA and TatB with each of the binding sites on TatC. **A** and **C.** TatA, TatB and TatC, assembled with TatB in the polar cluster site and TatA in its constitutive site for panels **B.** and **D.** TatA is in the polar cluster site and TatB in the constitutive site. Plots show the evolution of the minimum distances between L9 and either M205 or L206 during the 100 ns simulations. For both TatA and TatB, the simulations retain the close proximity between L9 and M205 at ~4 Å, while L9 to L206 is more variable, at around 8 Å.

**Figure S6.**
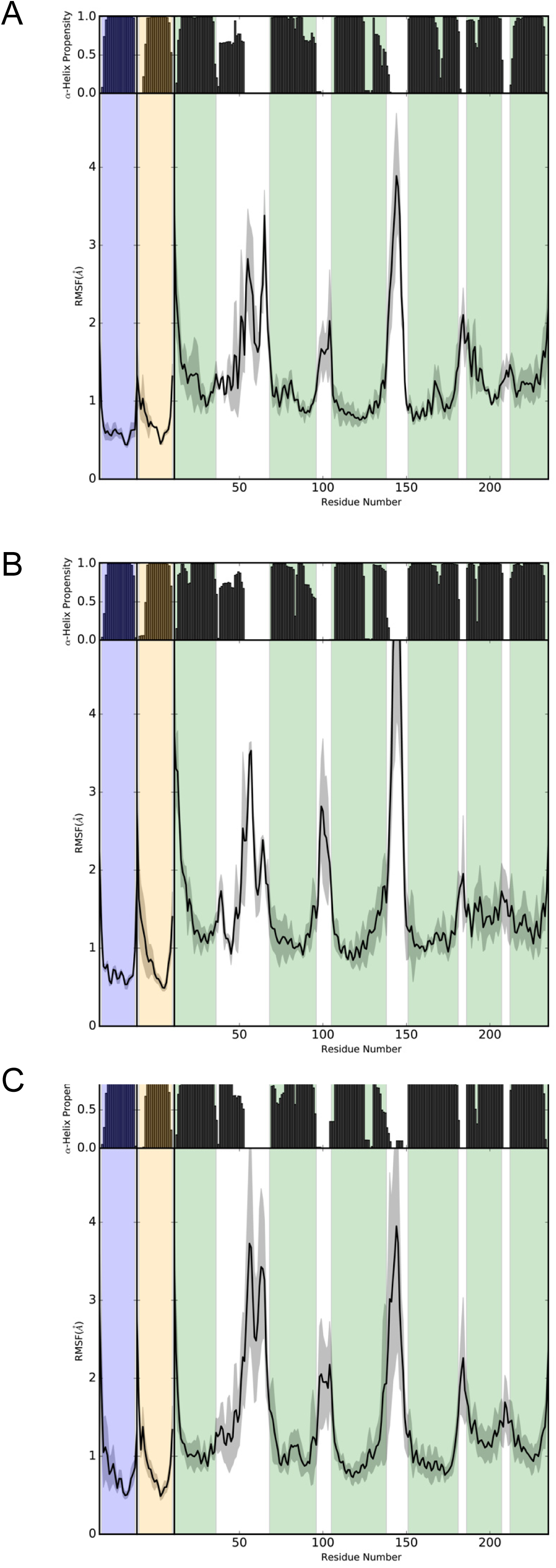
Structural stability plots for the modelled Tat protein complexes from molecular simulations. TatA, TatB and TatC, assembled with TatB in the polar cluster site and TatA in its constitutive site, for **A.** the heterotrimeric and **C.** heterononameric comples. For **B.** TatA is in the polar cluster site and TatB in its constitutive site in a heterotrimeric TatABC complex. The plots illustrate the retention of α-helical secondary structure (black bars) for each molecular system, with the lower panels in each plot showing the residue fluctuations. α-helical regions of the plots are coloured blue for TatA, yellow for TatB and green for TatC.

## Acknowledgements

This work was supported by the UK Biotechnology and Biological Sciences Research Council (through grants BB/N014545/1 and BB/L001306/1) and the UK Medical Research Council (through grant G1001640). We thank Professor Ben Berks and Dr Felicity Alcock for helpful discussion.

---

**Table S1.**
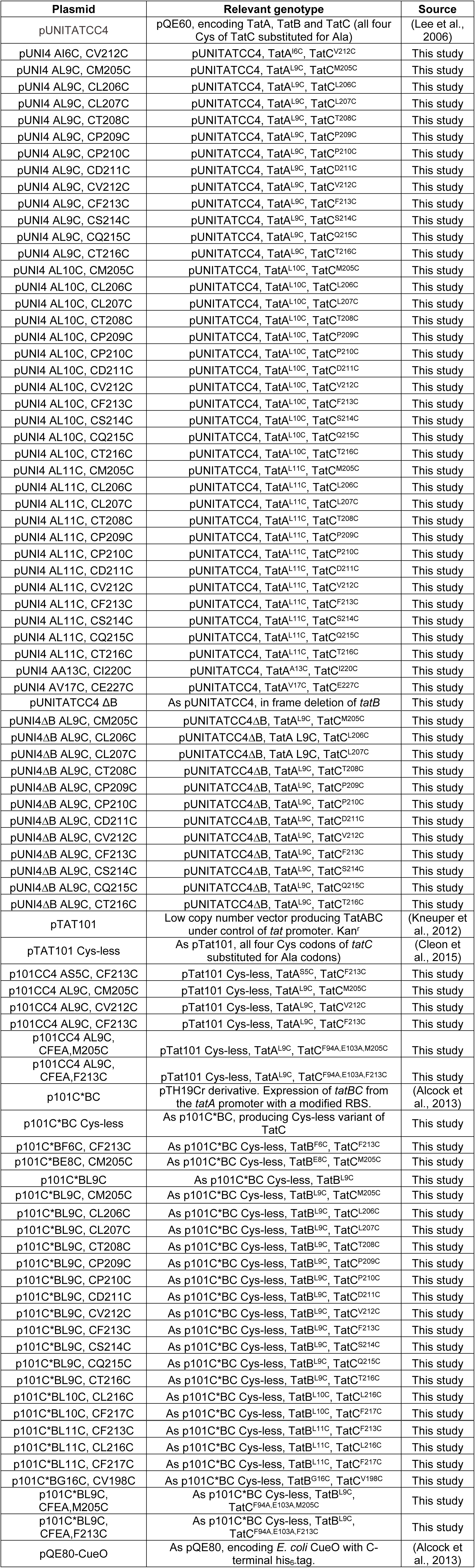
Plasmids used in this study

**Table S2.**
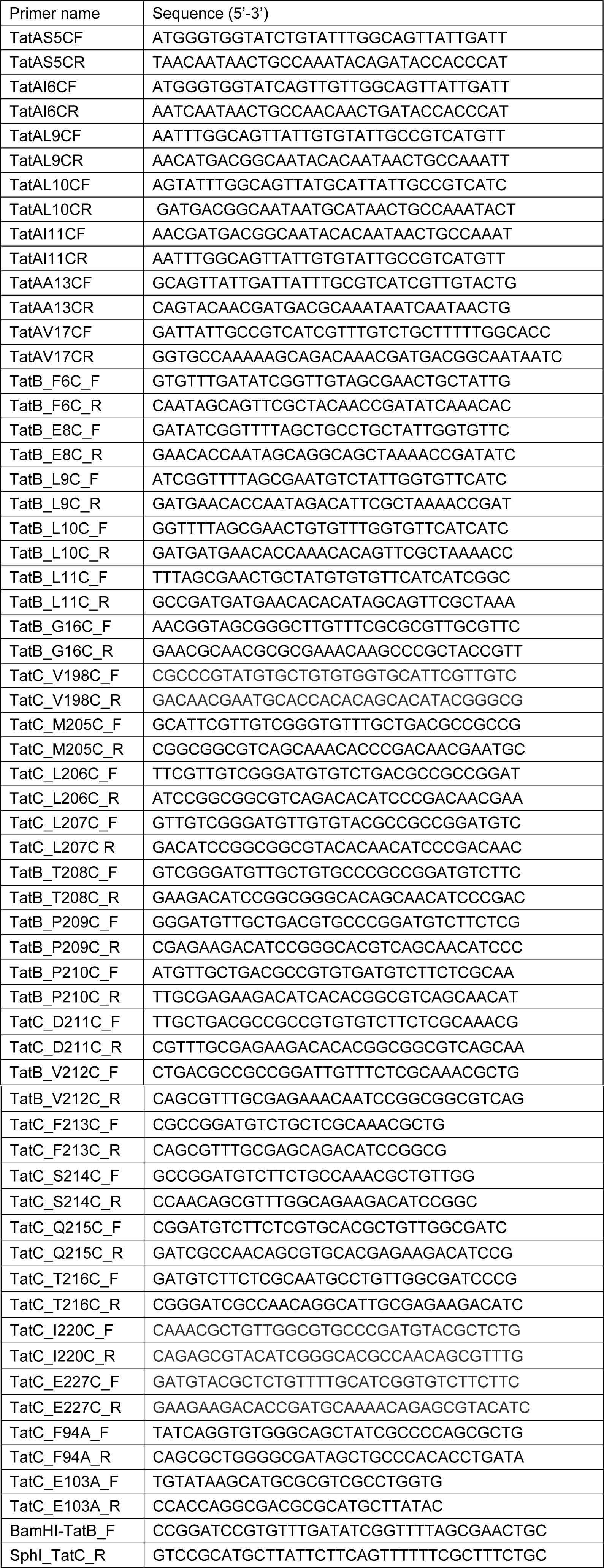
Oligonucleotides used in this study

## References

Alami, M., I. Luke, S. Deitermann, G. Eisner, H. G. Koch, J. Brunner, and M. Muller. 2003. Differential interactions between a twin-arginine signal peptide and its translocase in *Escherichia coli*. Mol. Cell. 12:937–946.

Alcock, F., M. A. Baker, N. P. Greene, T. Palmer, M. I. Wallace, and B. C. Berks. 2013. Live cell imaging shows reversible assembly of the TatA component of the twin-arginine protein transport system. Proc. Natl. Acad. Sci. of the USA. 110:E3650–3659.

Alcock, F., P. J. Stansfeld, H. Basit, J. Habersetzer, M. A. Baker, T. Palmer, M. I. Wallace, and B. C. Berks. 2016. Assembling the Tat protein translocase. eLife. 5.

Aldridge, C., X. Ma, F. Gerard, and K. Cline. 2014. Substrate-gated docking of pore subunit Tha4 in the TatC cavity initiates Tat translocase assembly. J. Cell. Biol. 205:51–65.

Aldridge, C., A. Storm, K. Cline, and C. Dabney-Smith. 2012. The chloroplast twin arginine transport (Tat) component, Tha4, undergoes conformational changes leading to Tat protein transport. J. Biol. Chem. 287:34752–34763.

Behrendt, J., and T. Bruser. 2014. The TatBC complex of the Tat protein translocase in *Escherichia coli* and its transition to the substrate-bound TatABC complex. Biochemistry. 53:2344–2354.

Behrendt, J., U. Lindenstrauss, and T. Bruser. 2007. The TatBC complex formation suppresses a modular TatB-multimerization in *Escherichia coli*. FEBS Lett. 581:4085–4090.

Berks, B.C. 2015. The twin-arginine protein translocation pathway. Annu. Rev. Biochem. 84:843–864.

Blummel, A.S., L. A. Haag, E. Eimer, M. Muller, and J. Frobel. 2015. Initial assembly steps of a translocase for folded proteins. Nature Comms. 6:7234.

Bolhuis, A., J. E. Mathers, J. D. Thomas, C. M. Barrett, and C. Robinson. 2001. TatB and TatC form a functional and structural unit of the twin-arginine translocase from *Escherichia coli*. J. Biol. Chem. 276:20213–20219.

Bussi, G., D. Donadio, and M. Parrinello. 2007. Canonical sampling through velocity rescaling. J Chem Phys.126:014101.

Casadaban, M.J., and S. N. Cohen. 1979. Lactose genes fused to exogenous promoters in one step using a Mu-lac bacteriophage: in vivo probe for transcriptional control sequences. Proc. Natl. Acad. Sci. of the USA. 76:4530–4533.

Cleon, F., J. Habersetzer, F. Alcock, H. Kneuper, P. J. Stansfeld, H. Basit, M. I. Wallace, B. C. Berks, and T. Palmer. 2015. The TatC component of the twin-arginine protein translocase functions as an obligate oligomer. Mol. Microbiol. 98:111–129.

Cline, K. 2015. Mechanistic Aspects of Folded Protein Transport by the Twin Arginine Translocase (Tat). J. Biol. Chem. 290:16530–16538.

Cline, K., and H. Mori. 2001. Thylakoid DeltapH-dependent precursor proteins bind to a cpTatC-Hcf106 complex before Tha4-dependent transport. J. Cell. Biol. 154:719–729.

Dabney-Smith, C., and K. Cline. 2009. Clustering of C-terminal stromal domains of Tha4 homo-oligomers during translocation by the Tat protein transport system. Mol. Biol. Cell. 20:2060–2069.

Dabney-Smith, C., H. Mori, and K. Cline. 2006. Oligomers of Tha4 organize at the thylakoid Tat translocase during protein transport. J. Biol. Chem. 281:5476–5483.

Darden, T., D. York, and L. Pedersen. 1993. Particle mesh Ewald: An Nlog(N) method for Ewald sums in large systems. J Chem Phys. 98:10089–10092.

de Jong, D.H., G. Singh, W. F. Bennett, C. Arnarez, T. A. Wassenaar, L. V. Schäfer, X. Periole, D. P. Tieleman, and S. J. Marrink. 2013. Improved Parameters for the Martini Coarse-Grained Protein Force Field. J Chem Theory Comput. 9:687–697.

Dominguez, C., R. Boelens, and A. M. Bonvin. 2003. HADDOCK: a protein-protein docking approach based on biochemical or biophysical information. J Am Chem Soc. 125:1731–1737.

Fritsch, M.J., M. Krehenbrink, M. J. Tarry, B. C. Berks, and T. Palmer. 2012. Processing by rhomboid protease is required for *Providencia stuartii* TatA to interact with TatC and to form functional homo-oligomeric complexes. Mol. Microbiol. 84:1108–1123.

Gerard, F., and K. Cline. 2007. The thylakoid proton gradient promotes an advanced stage of signal peptide binding deep within the Tat pathway receptor complex. J. Biol. Chem. 282:5263–5272.

Hess, B. H. Bekker, H.J. C. Berendsen, and J. G. E. M. Fraaije. 1997. LINCS: A Linear Constraint Solver for Molecular Simulations. J Comp Chem. 18:1463–1472.

Hu, Y., E. Zhao, H. Li, B. Xia, and C. Jin. 2010. Solution NMR structure of the TatA component of the twin-arginine protein transport system from gram-positive bacterium *Bacillus subtilis*. J Am Chem Soc. 132:15942–15944.

Huang, Q., F. Alcock, H. Kneuper, J. C. Deme, S. E. Rollauer, S. M. Lea, B. C. Berks, and T. Palmer. 2017. A signal sequence suppressor mutant that stabilizes an assembled state of the twin arginine translocase. Proc. Natl. Acad. Sci. USA. in press. doi: 10.1073/pnas.1615056114

Ize, B., N. R. Stanley, G. Buchanan, and T. Palmer. 2003. Role of the *Escherichia coli* Tat pathway in outer membrane integrity. Mol. Microbiol. 48:1183–1193.

Jack, R.L., F. Sargent, B. C. Berks, G. Sawers, and T. Palmer. 2001. Constitutive expression of *Escherichia coli tat* genes indicates an important role for the twin-arginine translocase during aerobic and anaerobic growth. J. Bacteriol. 183:1801–1804.

Jefferys, E., Z. A. Sands, J. Shi, M. S. Sansom, and P. W. Fowler. 2015. Alchembed: A Computational Method for Incorporating Multiple Proteins into Complex Lipid Geometries. J Chem Theory Comput. 11:2743–2754.

Kneuper, H., B. Maldonado, F. Jager, M. Krehenbrink, G. Buchanan, R. Keller, M. Muller, B. C. Berks, and T. Palmer. 2012. Molecular dissection of TatC defines critical regions essential for protein transport and a TatB-TatC contact site. Mol. Microbiol. 85:945–961.

Koch, S., M. J. Fritsch, G. Buchanan, and T. Palmer. 2012. The *Escherichia coli* TatA and TatB proteins have an N-out C-in topology in intact cells. J. Biol. Chem. 287:14420–14431.

Laemmli, U.K. 1970. Cleavage of structural proteins during the assembly of the head of bacteriophage T4. Nature. 227:680–685.

Leake, M.C., N. P. Greene, R. M. Godun, T. Granjon, G. Buchanan, S. Chen, R. M. Berry, T. Palmer, and B. C. Berks. 2008. Variable stoichiometry of the TatA component of the twin-arginine protein transport system observed by in vivo single-molecule imaging. Proc. Natl. Acad. Sci. of the USA. 105:15376–15381.

Lee, P.A., G. L. Orriss, G. Buchanan, N. P. Greene, P. J. Bond, C. Punginelli, R. L. Jack, M. S. Sansom, B. C. Berks, and T. Palmer. 2006. Cysteine-scanning mutagenesis and disulfide mapping studies of the conserved domain of the twin-arginine translocase TatB component. J. Biol. Chem. 281:34072–34085.

Lopilato, J., S. Bortner, and J. Beckwith. 1986. Mutations in a new chromosomal gene of *Escherichia coli* K-12, *pcnB,* reduce plasmid copy number of pBR322 and its derivatives. Mol Gen Genet. 205:285–290.

Ma, X., and K. Cline. 2010. Multiple precursor proteins bind individual Tat receptor complexes and are collectively transported. EMBO J. 29:1477–1488.

McDevitt, C.A., G. Buchanan, F. Sargent, T. Palmer, and B. C. Berks. 2006. Subunit composition and in vivo substrate-binding characteristics of *Escherichia coli* Tat protein complexes expressed at native levels. FEBS J. 273:5656–5668.

Mori, H., and K. Cline. 2002. A twin arginine signal peptide and the pH gradient trigger reversible assembly of the thylakoid [Delta]pH/Tat translocase. J. Cell. Biol. 157:205–210.

Oates, J., J. Mathers, D. Mangels, W. Kuhlbrandt, C. Robinson, and K. Model. 2003. Consensus structural features of purified bacterial TatABC complexes. J. Mol. Biol. 330:277–286.

Oostenbrink, C., A. Villa, A. E. Mark, and W.F. van Gunsteren. 2004. A biomolecular force field based on the free enthalpy of hydration and solvation: the GROMOS force-field parameter sets 53A5 and 53A6. J Comput Chem. 25:1656–1676.

Orriss, G.L., M. J. Tarry, B. Ize, F. Sargent, S. M. Lea, T. Palmer, and B. C. Berks. 2007. TatBC, TatB, and TatC form structurally autonomous units within the twin arginine protein transport system of *Escherichia coli*. FEBS Lett. 581:4091–4097.

Palmer, T., and B. C. Berks. 2012. The twin-arginine translocation (Tat) protein export pathway. Nat. Rev. Microbiol. 10:483–496.

Parrinello, M., and A. Rahman. 1981. Polymorphic transitions in single crystals: A new molecular dynamics method. J Applied Phys. 52:7182.

Pronk, S., S. Pall, R. Schulz, P. Larsson, P. Bjelkmar, R. Apostolov, M. R. Shirts, J. C. Smith, P. M. Kasson, D. van der Spoel, B. Hess, and E. Lindahl. 2013. GROMACS 4.5: a high-throughput and highly parallel open source molecular simulation toolkit. Bioinformatics. 29:845–854.

Ramasamy, S., R. Abrol, C. J. Suloway, and W. M. Clemons, Jr., 2013. The glove-like structure of the conserved membrane protein TatC provides insight into signal sequence recognition in twin-arginine translocation. Structure. 21:777–788.

Rodriguez, F., S. L. Rouse, C. E. Tait, J. Harmer, A. De Riso, C. R. Timmel, M. S. Sansom, B. C. Berks, and J. R. Schnell. 2013. Structural model for the protein-translocating element of the twin-arginine transport system. Proc. Natl. Acad. Sci. of the USA. 110:E1092-1101.

Rollauer, S.E., M. J. Tarry, J. E. Graham, M. Jaaskelainen, F. Jager, S. Johnson, M. Krehenbrink, S. M. Liu, M. J. Lukey, J. Marcoux, M. A. McDowell, F. Rodriguez, P. Roversi, P. J. Stansfeld, C. V. Robinson, M. S. Sansom, T. Palmer, M. Hogbom, B. C. Berks, and S. M. Lea. 2012. Structure of the TatC core of the twin-arginine protein transport system. Nature. 492:210–214.

Rose, P., J. Frobel, P. L. Graumann, and M. Muller. 2013. Substrate-dependent assembly of the Tat translocase as observed in live Escherichia coli cells. PLoS ONE. 8:e69488.

Sargent, F., U. Gohlke, E. De Leeuw, N. R. Stanley, T. Palmer, H. R. Saibil, and B. C. Berks. 2001. Purified components of the Escherichia coli Tat protein transport system form a double-layered ring structure. Eur. J. Biochem. /FEBS. 268:3361–3367.

Sargent, F., N. R. Stanley, B. C. Berks, and T. Palmer. 1999. Sec-independent protein translocation in *Escherichia coli.* A distinct and pivotal role for the TatB protein. J. Biol. Chem. 274:36073–36082.

Stansfeld, P.J., J. E. Goose, M. Caffrey, E. P. Carpenter, J. L. Parker, S. Newstead, and M. S. Sansom. 2015. MemProtMD: Automated Insertion of Membrane Protein Structures into Explicit Lipid Membranes. Structure. 23:1350–1361.

Stansfeld, P.J., and M. S. Sansom. 2011. From Coarse Grained to Atomistic: A Serial Multiscale Approach to Membrane Protein Simulations. J Chem Theory Comput. 7:1157–1166.

Tarry, M.J., E. Schafer, S. Chen, G. Buchanan, N. P. Greene, S. M. Lea, T. Palmer, H. R. Saibil, and B. C. Berks. 2009. Structural analysis of substrate binding by the TatBC component of the twin-arginine protein transport system. Proc. Natl. Acad. Sci. of the USA. 106:13284–13289.

Wexler, M., F. Sargent, R. L. Jack, N. R. Stanley, E. G. Bogsch, C. Robinson, B. C. Berks, and T. Palmer. 2000. TatD is a cytoplasmic protein with DNase activity. No requirement for TatD family proteins in sec-independent protein export. J. Biol. Chem. 275:16717–16722.

Zhang, Y., Y. Hu, H. Li, and C. Jin. 2014a. Structural basis for TatA oligomerization: an NMR study of *Escherichia coli* TatA dimeric structure. PLoS ONE. 9:e103157.

Zhang, Y., L. Wang, Y. Hu, and C. Jin. 2014b. Solution structure of the TatB component of the twin-arginine translocation system. Biochim. Biophys. Acta. 1838:1881–1888.

Zoufaly, S., J. Frobel, P. Rose, T. Flecken, C. Maurer, M. Moser, and M. Muller. 2012. Mapping precursor-binding site on TatC subunit of twin arginine-specific protein translocase by site-specific photo cross-linking. J. Biol. Chem. 287:13430–13441.

